# Adhesion-mediated mechanosignaling forces mitohormesis

**DOI:** 10.1101/2020.03.06.979583

**Authors:** Kevin M. Tharp, Ryo Higuchi-Sanabria, Greg A. Timblin, Breanna Ford, Carlos Garzon-Coral, Catherine Schneider, Jonathon M. Muncie, Connor Stashko, Joseph R. Daniele, Andrew S. Moore, Phillip A. Frankino, Sagar S. Manoli, Hao Shao, Alicia L. Richards, Kuei-Ho Chen, Gregory M. Ku, Marc Hellerstein, Daniel K. Nomura, Karou Saijo, Jason Gestwicki, Alexander R. Dunn, Nevan J. Krogan, Danielle L. Swaney, Andrew Dillin, Valerie M. Weaver

## Abstract

Mitochondria control eukaryotic cell fate by producing the energy needed to support life and the signals required to execute programmed cell death. The biochemical milieu is known to affect mitochondrial function and contribute to the dysfunctional mitochondrial phenotypes implicated in cancer and the morbidities of ageing. However, the physical characteristics of the extracellular matrix are also altered in cancer and in aging tissues. We demonstrate that cells sense the physical properties of the extracellular matrix and activate a mitochondrial stress response that adaptively tunes mitochondrial function *via* SLC9A1-dependent ion exchange and HSF1-dependent transcription. Overall, our data indicate that adhesion-mediated mechanosignaling may play an unappreciated role in the altered mitochondrial functions observed in aging and cancer.

**Graphical Abstract:** 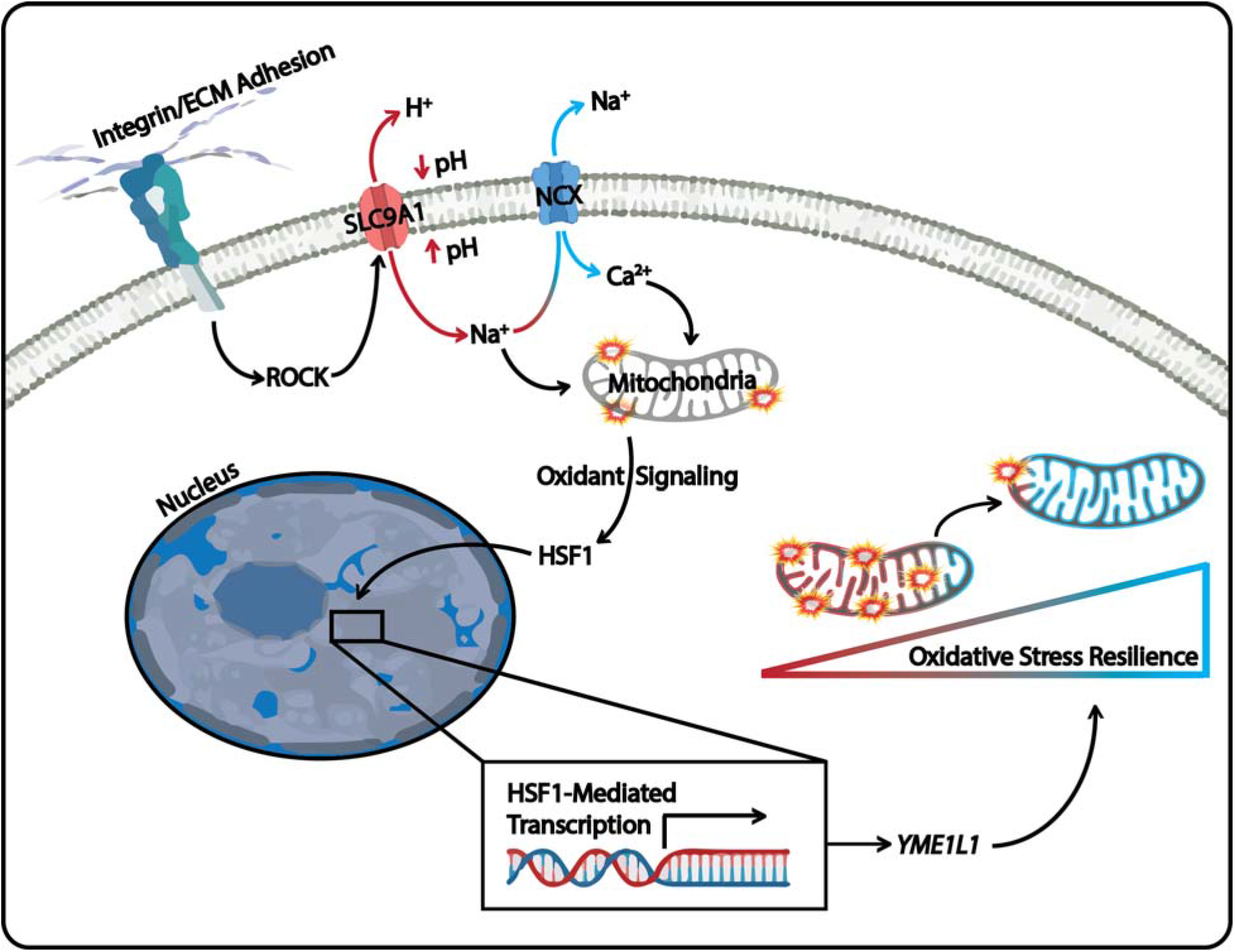

---

Alterations in mitochondrial function are essential for the ability of cancer cells to rapidly proliferate and metastasize and aging cells to regulate senescence phenotypes induced by DNA damage. Mitochondria provide a privileged metabolic compartment where oxidative phosphorylation (OxPhos) consumes oxygen and reducing equivalents to produce ATP. While OxPhos provides an efficient means to produce ATP, it can create collateral cellular damage through the release of reactive oxygen species (ROS), which can oxidize proteins, lipids, and nucleic acids. In response to elevated mitochondrial ROS exposure, cancer and aging cells activate adaptive stress responses that allow them to harness ROS-mediated proliferation and migration effects without activating ROS-mediated cell death (*1–4*). These types of oxidative stress resilience (OxSR) programs alter cellular metabolism to enhance ROS buffering in the cytosol to limit damage caused by mitochondrial ROS.

The overproduction of ROS *via* mitochondrial dysfunction is thought to occur because of the biochemical composition of the aged or tumor microenvironment (TME) (*5–8*). However, the physical characteristics of the cellular microenvironment are also altered by cancer or aging - and can affect cell fate, function, and metabolism through the activity of stress- and ROS-associated transcription factors (*5, 9–11*). Sensing of the mechanical properties of the extracellular matrix (ECM) can also affect cellular metabolism by regulating the levels and/or activity of cytoplasmic enzymes responsive to mechanosignaling-induced cytoskeletal dynamics (Papalazarou et al., 2020; Park et al., 2020). Cellular mechanosignaling relies on adhesion receptors, such as integrins, transducing signals that mechanically entrain the cytoskeleton to the ECM, and these cytoskeletal remodeling events can affect the topological distribution of metabolic organelles, cargoes, and enzymes (*11–13*). Cytoskeletal dynamics also play a critical role in the regulation of mitochondrial structure (Helle et al., 2017; Manor et al., 2015; Moore et al., 2016) and mechanosensitive transcription factors can alter mitochondrial gene expression (*14*). Because mitochondrial structure influences mitochondrial function, we sought to determine if and how adhesion-mediated mechanosignaling affects mitochondrial function (Figure 1A).

**Figure 1.**
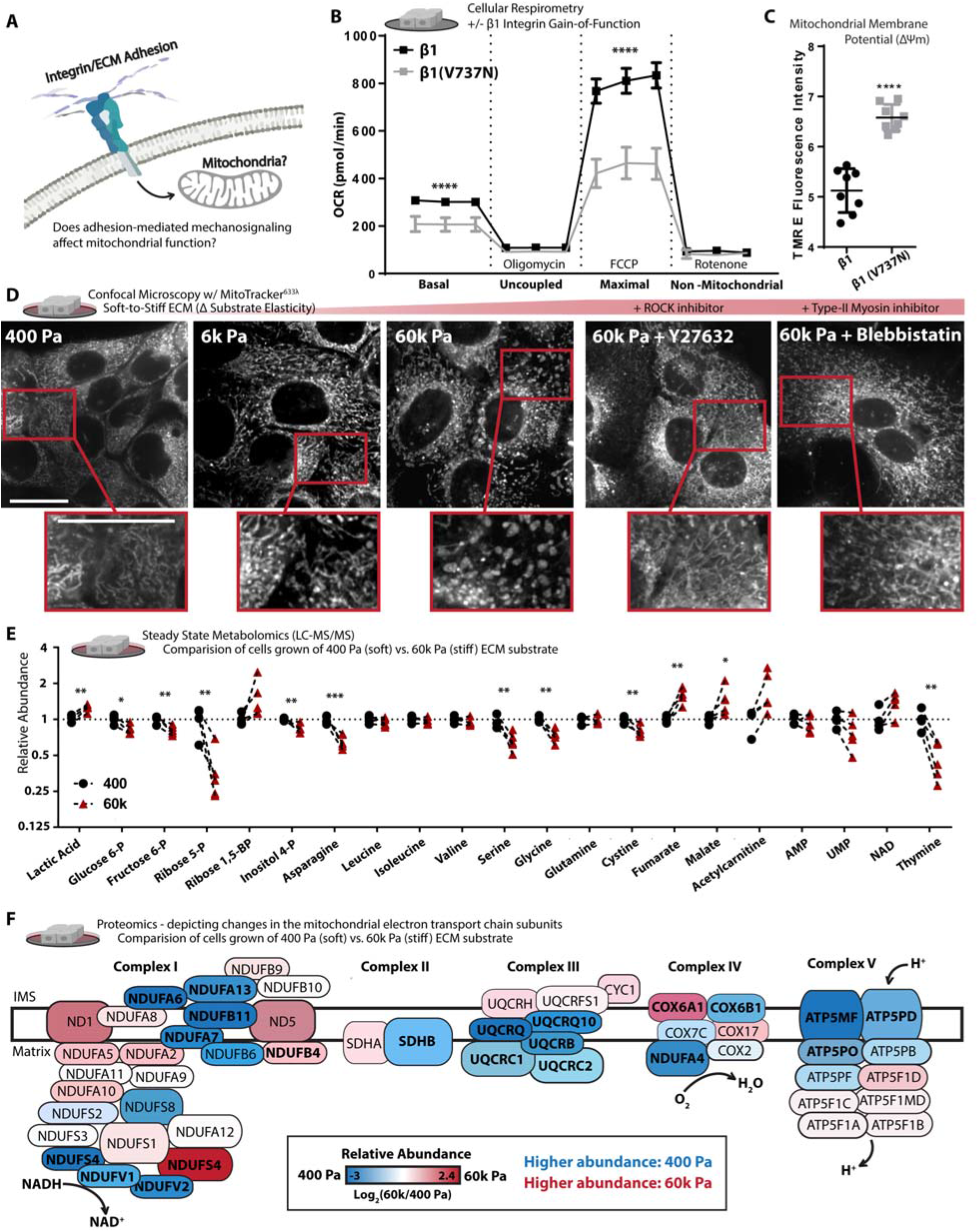
Adhesion-mediated mechanosignaling alters mitochondrial structure and function of human mammary epithelial cells (MECs). A. Graphical representation of the experimental question. B. Mitochondrial oxygen consumption rate (OCR) of β1-integrin or β1(V7373N) expressing cells (100k cells per well, n=5 wells, 3 replicate measures, repeated 3 times), mitochondrial stress test conditions: uncoupled = oligomycin [1 μM], maximal = Trifluoromethoxy carbonylcyanide phenylhydrazone (FCCP) [1 μM], non-mitochondrial = antimycin A [1 μM] and rotenone [1 μM]. C. Mitochondrial membrane potential, measured after 1 h treatment of Tetramethylrhodamine ethyl-ester (TMRE) [10 nM] (n = 2 wells, repeated 4 times). D. Confocal Microscopy depicting mitochondrial network structure in PFA-fixed cells cultured on varied soft-to-stiff fibronectin coated [6 μM/cm2] polyacrylamide hydrogels (soft-to-stiff ECM), for 24 h +/- y27632 [10 μM] or Blebbistatin [10 μM], stained with mitotracker (deep red FM) [100 nM]. E. Selection of metabolites measured with (LC-MS/MS) from MECs cultured on soft of stiff ECM for 24 h, fold change relative to 400 Pa. (n=4-5 biological replicates LC-MS/MS run together, repeated 2 times) F. Relative abundance (fold change) of mitochondrial ETC subunits measured *via* timsTOF LC-MS of MECs cultured on soft or stiff ECM for 24 h (n=3 biological replicates). Bolded text indicates *P of < 0.05 or less, locations and sizes of ETC subunits graphically depicted are approximate and not to molecular scale. *Data shown represents ± SEM. *P < 0.05, **P < 0.01, ***P < 0.005, < 0.0001 via twotailed unpaired Student t test*.

## Results

### Mechanosignaling alters mitochondrial structure and function

We investigated the relationship between adhesion-dependent mechanosignaling and mitochondrial function by exogenously expressing a β1-integrin “gain of function” mechanosignaling model in nonmalignant human mammary epithelial cells (MECs; MCF10A). Expression of the β1-integrin (V737N, point mutation) promotes focal adhesion assembly, phosphorylation of focal adhesion kinase (FAK) (Figure S1A), cytoskeletal remodeling, actomyosin tension (*15*), and the suppression of mitochondrial oxygen consumption (Figure 1B). Cellular respiration is primarily a product of OxPhos, in which the mitochondrial electron transport chain (ETC) consumes oxygen and reducing equivalents to produce ATP. Contrary to the paradigm that suppressed mitochondrial function (e.g. reduced respiration) occurs due to the loss of the mitochondrial membrane potential (ΔΨm), the β1(V737N)-integrin expressing MECs had higher, not lower, mitochondrial membrane potential (Figure 1C). To ensure that these phenotypes were not due to an indirect effect of β1(V737N)-integrin expression, we varied the surface density coating of fibronectin, an ECM component and integrin adhesion ligand that increases mechanosignaling *via* β1-integrin (*16*). MECs plated on a higher density of a fibronectin surface coating [60 μM/cm^2^], also showed repression of mitochondrial oxygen consumption, similar to β1(V737N)-integrin expression (Figure S1B). Furthermore, activating integrin mechanosignaling with acute exposure to manganese (Mn^2+^) (*17*) also suppressed mitochondrial oxygen consumption (Figure S1C). The data indicate that that increased integrin mechanosignaling impacts mitochondrial function.

Since integrin mechanosignaling is highly sensitive to the stiffness of the adhesion substrate, which affects mitochondrial respiration (oxygen consumption), we examined the mitochondrial morphology of MECs cultured for 24 hours on fibronectin coated [6 μM/cm^2^] polyacrylamide hydrogel gels ranging in elasticity (stiffness) between normal breast (400 Pa) and tumor (6k – 60k Pa) ECM (*18, 19*) (note: tissue culture polystyrene is supraphysiological, ~3G Pa). MECs cultured on this range of biologically relevant ECM elasticities displayed a variety of mitochondrial morphologies, ranging from thin interconnected filaments (400 Pa), to thickened filaments (6k Pa), and then ~300 nM diameter fragments with toroidal shapes (60k Pa) (Figure 1D and S1D). Cells respond to ECM stiffness by ligating ECM adhesion receptors that induce Rho-GTPase and Rho-associated protein kinase (ROCK) cytoskeletal remodeling and increase actomyosin tension *via* type-II myosins (*20*). We therefore bypassed the adhesion receptor ligation step and induced downstream mechanosignaling *via* an inducible ROCK, ROCK::ER (*21*), which was sufficient to suppress mitochondrial oxygen consumption (Figure S1E & S1A). In contrast, pharmacological inhibition of ROCK with Y27632 or type-II myosins with blebbistatin reduced the prevalence of the thick or toroidal mitochondrial fragments in MECs plated on the stiff ECM (Figure 1D), and restored mitochondrial oxygen consumption in the ROCK::ER cells (Figure S1F). Finally, β1(V737N)-integrin expressing cells displayed a fragmented/toroidal mitochondrial morphology, even on the soft ECM (Figure S1G) in direct contrast to MECs expressing a wild-type β1 integrin. The data indicate that mitochondrial structure and function is sensitive to the stiffness of the ECM through integrin- and ROCK-mediated mechanosignaling.

Because mitochondrial function affects many aspects of cellular metabolism, we broadly examined the steady state levels of polar metabolites present in MECs cultured on ECM surfaces that mimic the soft normal ECM (400 Pa) or stiff tumor ECM (60k Pa) (Figure 1E and Video S1-5). MECs cultured on the stiff ECM possessed higher levels of lactate and lower levels of upstream glycolytic or pentose phosphate pathway (PPP) intermediates, which may indicate increased flux through those pathways. We also noted lower levels of serine (*22*) and increased levels of tricarboxylic acid cycle (TCA cycle) intermediates such as malate and fumarate, an oncometabolite (*23*), which could indicate that TCA cycle flux has reduced. Indeed, previous studies have indicated that TCA cycle impairment can affect mitochondrial structure (*24*). Since TCA cycle flux is largely dependent on the activity of the ETC, we mapped the compositional changes in ETC subunit abundance with mass spectrometry based proteomics (Figure 1F). We found that a number of critical ETC subunits changed in abundance due to ECM stiffness and their relative levels can alter properties of mitochondrial function (*25*). These mechanosignaling induced compositional changes in the ETC could explain the reduction of mitochondrial oxygen consumption, and the increased ΔΨm observed when integrin mechanosignaling is high due to decreased entry of electrons via NADH (Complex I) and proton pumping from the inner membrane space (IMS) into the mitochondrial matrix (MM) through ATP Synthase (Complex V).

### Hyperglycemia and stiff ECM facilitate similar mitochondrial responses

Hyperglycemia [>5 mM glucose] is a biochemical stress that induces mitochondrial fragmentation, raises intracellular pH (pH_i_) (*26*), lowers extracellular pH (pH_e_), and increases mitochondrial membrane potential in cultured cells (*27*). To explore whether the fragmented/toroidal mitochondrial morphologies induced by high cytoskeletal tension, were similar to those induced by hyperglycemia, we increased the media glucose concentration from 5 mM (“low glucose”) to 25 mM (“high glucose”) for MECs plated on the soft ECM (400 Pa) and examined changes in mitochondrial organization. Lattice light sheet microscopy (LLSM), which permits live cell imaging with limited phototoxicity (*28*), revealed that exposing MECs to hyperglycemia induced a rapid transition of mitochondrial morphology from a filamentous network into fragmented/toroidal structures (Figure S1H and Video 5-6), comparable to MECs cultured on stiff ECM (Figure 1D & S1D). Moreover, cells exposed to hyperglycemia or plated on stiff ECM express similar gene profiles that have been implicated in the mitochondrial unfolded protein response (UPR^mt^) (*29, 30*) (Figure S1I). Since both hyperglycemia and stiff ECM induce these mitochondrial stress response genes, we hypothesized that the reorganization of the mitochondria may reflect a pro-survival stress response (*31, 32*).

Because ECM stiffness induces mitochondrial reorganization by inducing cytoskeletal tension (Figure 1D), and the similarities between hyperglycemia and ECM stiffness, we asked if hyperglycemia was sufficient to increase ECM stiffness. Atomic force microscopy (AFM) indentation revealed that hyperglycemia significantly enhanced cortical tension in MECs (Figure S1J). These findings were confirmed in a second cell line, the MDA-MB-231 MECs, which is a model of triple negative human breast cancer. Finally, similar to the reduced respiration rate induced by manipulating cytoskeletal tension, hyperglycemia also reduced mitochondrial oxygen consumption (Figure S1K). These findings demonstrate that biochemical and physical cues appear to stimulate similar changes in mitochondrial structure and function, and these changes may occur through the same stress response.

Mitochondrial fragmentation is thought to coordinate with mitophagy (autophagosome-mediated degradation of mitochondria) to repair dysfunctional mitochondria that have reduced mitochondrial membrane potential or increased reactive oxygen species (ROS) production/leak (*33–35*). However, our data indicated that the mitochondrial fragmentation induced by stiff ECM had elevated mitochondrial membrane potential (Figure 2B). Thus, we reasoned that large mitochondrial fragments with toroidal morphologies likely arise through a different mechanism than has been previously described. Mitochondrial membrane potential reflects a pH differential between the mitochondrial inner membrane space and the mitochondrial matrix, but it’s measurement with lipophilic cations (TMRE) can be sensitive to changes in intracellular pH (pH_i_) and calcium (Ca^2+^) (*36*). To verify that cytosolic pH changes were not confounding our measurement of ΔΨm, we measured pH_i_ of MECs cultured on the stiff ECM or exposed to hyperglycemia and found that both stresses increased pHi (Figure 2C). ROCK, a key mechanosignaling kinase, regulates the activity of SLC9A1 (Na^+^/H^+^ Exchanger 1 (NHE1)), which functions to increase the pH of the adhesion-proximal cytosol to facilitate conformational changes in FAK which are critical for its phosphorylation and mechanosignaling downstream of integrin adhesions (*37, 38*). Since both hyperglycemia and ROCK activity increase pH_i_ (*26, 39*), we prevented the coordinated decrease of pH_e_ (extracellular acidification, H^+^ pumped out of the cell) induced by hyperglycemia *via* inhibition of ROCK or SLC9A1 (Figure 2D).

**Figure 2.**
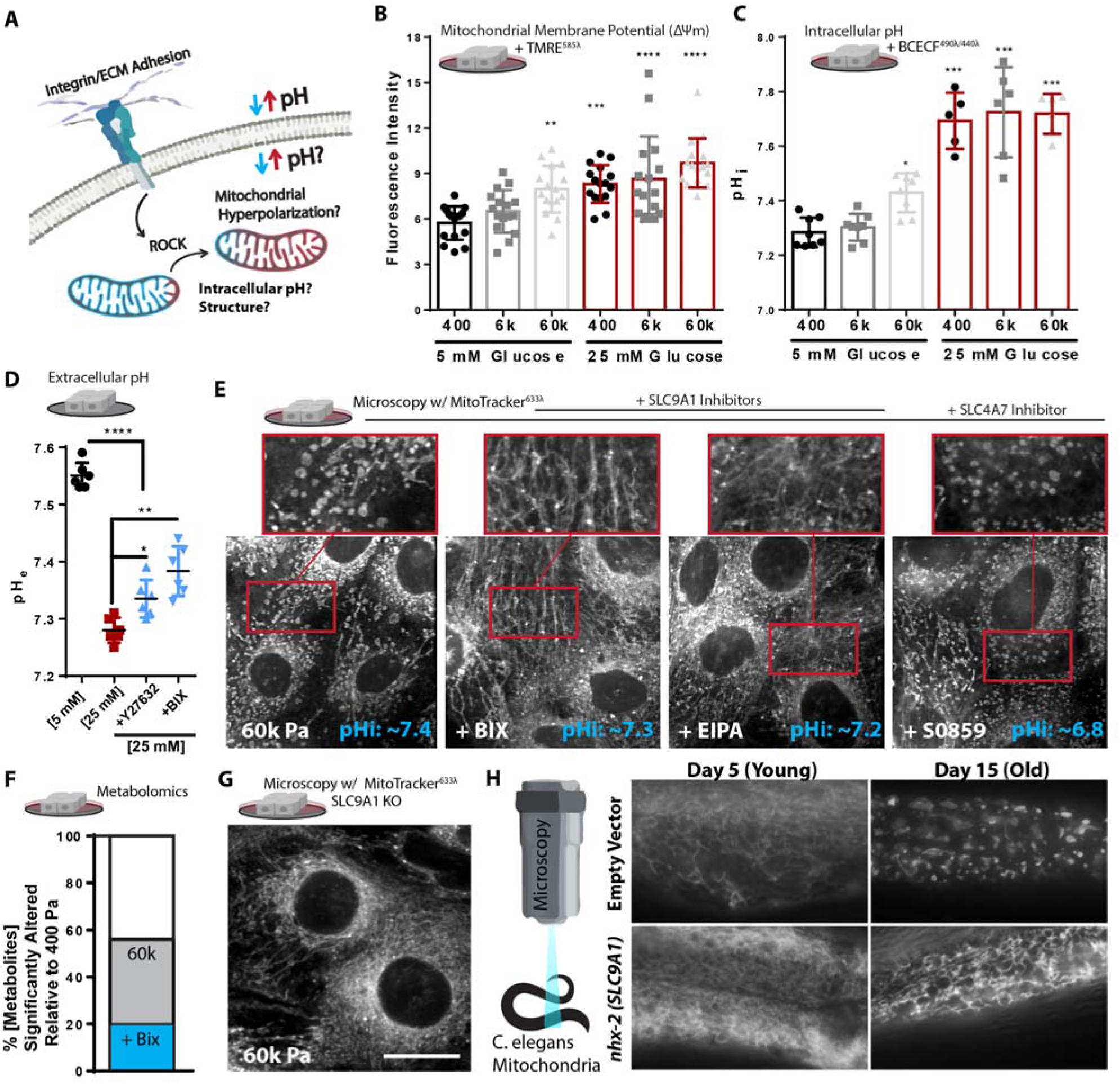
SLC9A1 facilitates stiff ECM induced mitochondrial programming. A. Graphical representation of the experimental question, because TMRE intensity (Figure 1C) could be an artifact of intracellular pH. B. Mitochondrial membrane potential, measured after 1 h treatment of Tetramethylrhodamine ethyl-ester (TMRE) [10 nM] (n = 4 replicate PA gels repeated 4 times, shown together). C. Intracellular pH (pHi) of cells grown of varied stiffness PA gels +/- glucose [25 mM], measured via 2’,7’-Bis-(2-Carboxyethyl)-5-(and-6)-Carboxyfluorescein, Acetoxymethyl Ester (BCECF) [1 μM] (n=2 replicate PA gels repeated 3-4 times, shown together). D. pH of media surrounding MCF10A cells cultured in glucose [5 or 25 mM] +/- DMSO, Y27632 [10 μM], or BIX [500 nM] with 200k cells/well of 12 well plate after 48 h (n=3 repeated 2 times, shown together). E. Confocal microscopy depicting mitochondrial network structure and caption depicting of pHi measurements (mean of n=5) in MECs on 60k Pa surfaces treated with BIX [500 nM], EIPA [10 μM], S0859 [50 μM], or vehicle for 24 h, mitotracker (deep red FM) [100 nM] and BCECF [1 μM] (n=2 replicate PA gels repeated 3 times). F. Metabolomics (LC-MS/MS) of cells cultured on 400 or 60k ECM for 24 h, % significantly altered relative to 400 Pa +/- BIX [500 nM]. (n=4-5 biological replicates LC-MS/MS run together, repeated 2 times) G. Confocal microscopy depicting mitochondrial network structure of *SLC9A1* KO cells on 60k Pa surfaces for 24 h, mitotracker (deep red FM) [100 nM]. H. Representative microscopy depicting mitochondrial network structure of live *C. elegans* expressing MLS::mRuby (mitochondrial matrix) grown on empty vector or *nhx-2 (SLC9A1* orthologue) RNAi from hatch of 5 d or 15 d old animals. *Data shown represents ± SEM. *P < 0.05, **P < 0.01, ***P < 0.005, < 0.0001 via oneway ANOVA with Tukey test for multiple comparisons to 400 Pa 5 mM Glucose (B and C) or 25 mM glucose (D*).

### SLC9A1-mediated ion exchange affects mitochondrial structure and function

To test if the elevated pH_i_ was responsible for the altered mitochondrial morphology observed in MECs cultured on stiff ECM, we lowered pH_i_ by inhibiting SLC9A1 with BIX and EIPA, and SLC4A7 (Na^+^/HCO^3-^ cotransporter) with S0859. While all of these interventions lowered pHi in MECs on stiff ECM to levels equivalent or lower than soft ECM, only SLC9A1 inhibition restored the filamentous mitochondrial morphology (Figure 2E & S2A). These data suggest that a pHi-independent effect of SLC9A1 may be responsible for the mitochondrial morphology induced by stiff ECM. SLC9A1 inhibition also restored the concentrations of approximately 60% of the significantly altered polar metabolites we measured in MECs plated on the stiff ECM back to the concentrations observed in MECs on soft ECM (Figure 2F). SLC9A1 inhibition also rescued the impaired mitochondrial oxygen consumption caused by β1(V737N)-integrin or ROCK::ER expression (Figure S2B-D). CRISPR-mediated knockout of SLC9A1 in MECs (*SLC9A1-KO*) resulted in MECs that maintained a filamentous mitochondrial morphology on stiff ECM (Figure 2G). *SLC9A1-KO* was sufficient to normalize mitochondrial respiration in MECs exposed to hyperglycemia, as well as those on high-density fibronectin coating (Figure S2F-G). To explore the physiological relevance of these findings, we examined the ability of SLC9A1 to affect mitochondrial morphology in *C. elegans*, a model organism that is amenable to live microscopy of mitochondria and genetic manipulations (*40*) that has been used extensively to study OxSR (*41*). RNAi-mediated knockdown of *nhx-2* (*SLC9A1* orthologue) prevented the mitochondrial fragmentation/toroidal phenotype typically found in the aged gut epithelium of this organism (Figure 2H).

As a byproduct of H^+^ efflux (raising pH_i_, lowering pH_e_) SLC9A1 facilitates Na^+^ import that subsequently reverses the directionality of the Na^+^/Ca^2+^ exchangers (NCX, SLC8A1-3), a process which ultimately causes mitochondrial ROS production *via* mitochondrial calcium (Ca^2+^) overload (*42, 43*). Additionally, cellular Na^+^ can affect the solubility of mitochondrial Ca^2+^ phosphate precipitates, fluidity of the mitochondrial inner membrane, and ROS production from Complex III of the ETC (*44*). To determine if adhesion-dependent cytoskeletal tension regulates mitochondrial Ca^2+^ content through SLC9A1 activity, we measured mitochondrial and total cellular Ca^2+^ levels in MECs plated on a range of soft-to-stiff ECM. Imaging of Rhod2-AM and Calcium Green-1 AM revealed that mitochondrial Ca^2+^ concentration was highest on the stiff ECM, and could be reduced either by inhibiting or knocking out *SLC9A1* (Figure 3B). SLC9A1 inhibition was also able to suppress the mitochondrial ROS production induced by mitochondrial Ca^2+^ loading in the CGP37157-treated cells (Figure 3C & S3A-B). Taken together, these data suggest that SLC9A1 alters mitochondrial function and ROS production.

**Figure 3.**
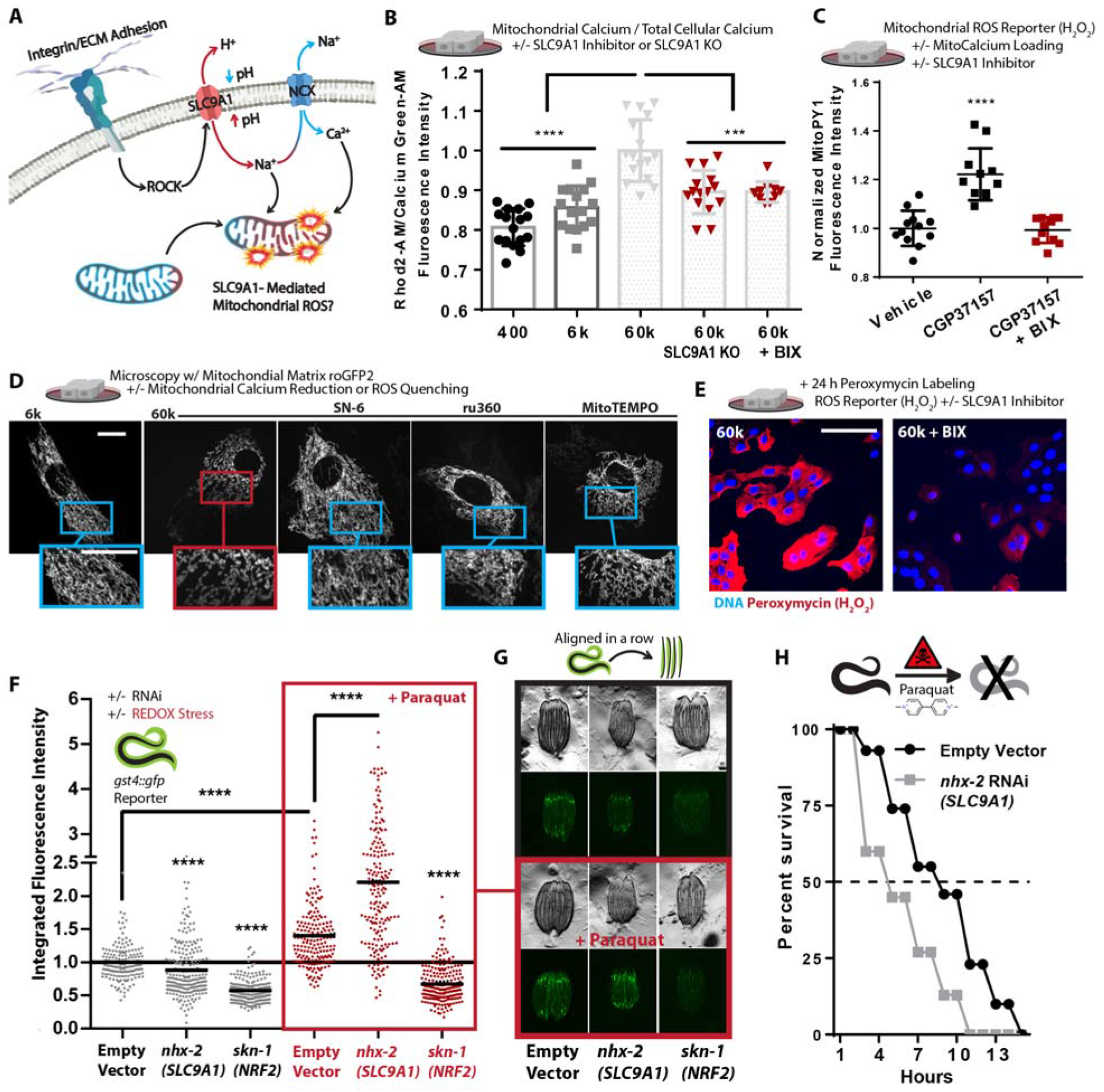
SLC9A1 facilitates mitochondrial oxidative stress. A. Graphical schematic indicating how SLC9A1 affects mitochondrial oxidative stress. B. Calcium content of MCF10A cells cultured on soft-to-stiff ECM for 24 h, treated with Rhod2-AM [2 μM] (mitochondrial) and Calcium Green-1-AM [2 μM] (intracellular). (n=4 replicates, repeated 4 times). C. Mitochondrial H_2_O_2_ production of cells cultured on 6k Pa surfaces and treated with BIX [500 nM] or vehicle for 24 h and then MitoPy1 [5 μM] and vehicle or CGP37157 [1 μM] for 1 h (n=6, repeated 2 times). D. Confocal microscopy depicting mitochondrial network structure of PFA fixed MECs on 6k or 60k Pa ECM treated with 1 μM ru360 (MCU inhibitor), 10 μM SN-6 (NCX reverse mode inhibitor, opposite direction of CCP37157), and 2 μM MitoTEMPO for 24 h. (Scale Bar: 10 μm) E. Confocal microscopy of peroxymycin (H_2_O_2_) (*22*) staining over 24 h on 60k Pa ECM +/- BIX [500 nM], quantitated in Figure S3I. F-G. *gst-4p::gfp* reporter fluorescent intensity of *C.elegans* measured with a large particle cytometer, +/- paraquat [50 mM] (n=177, 206, 190, 187, 191, and 215 animals in order left to right) with representative images (G) of *C. elegans* quantified, repeated 3 times. H. *C. elegans* survival in 50 mM paraquat at 1 d; animals grown from hatch on *nhx-2* RNAi vs empty-vector control (80 worms per condition, repeated 3 times). *Data shown represents ± SEM. *P < 0.05, **P < 0.01, ***P < 0.005, < 0.0001 via twotailed unpaired Student t test in (D) and one-way ANOVA with Tukey test for multiple comparisons in (B and C*).

Increased fibronectin surface density (Figure S3C-D), hyperglycemia (Figure S3E), and Mn^2+^ treatment also increased mitochondrial Ca^2+^ and ROS production, in part through the activity of SLC9A1 (Figure S3G). Consistently, treating cells with CGP37157 or kaempferol (mitochondrial calcium uniporter (MCU) activator), to increase mitochondrial Ca^2+^ content, was sufficient to induce the fragmented/toroidal mitochondrial morphology in MECs plated on the soft ECM (Figure S3H). Suppression of mitochondrial ROS, with 2-(2,2,6,6-tetramethylpiperidin-1-oxyl-4-ylamino)-2-oxoethyl)-triphenylphosphonium chloride (MitoTEMPO), or suppression of the mitochondrial Ca^2+^ loading *via* selective inhibition of the reverse-mode of NCX exchangers, with SN-6, and the MCU, with ru360, were also sufficient to prevent the fragment/toroid formation on stiff ECM (Figure 3D and Video S8-9), providing additional evidence that Ca^2+^ overload and ROS were causative of mitochondrial remodeling.

To directly test whether ECM stiffness could impact ROS production, we next monitored the ROS production in MECs on a range of soft-to-stiff ECM over the course of 24 hours (*22*). As expected, we found that cells seeded on stiff ECM produced more ROS than those on soft ECM, in an SLC9A1 dependent fashion (Figure 3E & S3I). Additionally, ROS production in ROCK:ER cells was also suppressed by SLC9A1 inhibition (Figure S3J). Finally, to determine whether the functional role of SLC9A1 on ROS-mediated oxidative stress was conserved in a whole organism, we tested its impact on oxidative stress in *C. elegans*. Using an reporter for oxidative stress (*gst-4p::gfp*), we found that *nhx-2* (*SLC9A1*) knockdown reduced the expression of the reporter, indicating a lower basal level of ROS production/response in these animals (Figure 3F-G). In agreement with the free radical theory of ageing, which postulates that lifespan is shortened due to accumulated oxidation-mediated damage, we found that the lifespan of the *C. elegans nhx-2* knockdown was dramatically extended (Figure S3K).

Since oxidative stress primarily regulates the expression of *gst-4p* through the transcriptional activity of *skn-1* (*NRF-2* orthologue), which activates the expression of genes with antioxidant response elements (ARE) in their promoters (e.g. *gst-4p*), we used a knockdown of *skn-1/NRF-2* as a negative control. As a positive control, we treated the *gst-4p::gfp* reporter animals with paraquat, an herbicide that promotes mitochondrial ROS leak/production (*45*), which increased oxidative stress reporter activity. Surprisingly, paraquat treatment induced a more robust *gst-4p::gfp* reporter response to paraquat in *nhx-2* knockdown animals (Figure 3F-G). This result suggests that *nhx-2* knockdown exacerbated paraquat-induced mitochondrial oxidative stress, likely because OxPhos could not be throttled (Figure 2D-F & S2A-F). To test the difference in oxidative stress sensitivity in *nhx-2* knockdown animals, we assayed the survival of these animals in response to paraquat-induced oxidative stress. Corroborating the *gst-4p::gfp* reporter measurements (Figure 3F-G), the *nhx-2* knockdown animals were more sensitive to paraquat exposure than control animals (Figure 3H). Accordingly, these results indicate that SLC9A1-activity induces OxSR *via* sub-lethal mitochondrial ROS, which promotes mitochondrial reorganization (toroid/fragment formation).

The greater induction of *gst-4p::gfp* and the rapidity of death observed in the *nhx-2* knockdown animals suggests they were less adapted to manage the ROS-mediated oxidative stress induced by paraquat. Since greater SLC9A1 activity appeared to promote mitochondrial ROS production (Figure 3C, 3E, S3G, & S3I-J) we hypothesized that because the *nhx-2* knockdown animals experienced lower basal levels of ROS exposure, they were not pre-adapted to survive the paraquat exposure. It has been reported that mitochondrial stresses, particularly ROS production and respiratory dysfunction, promote adaptive reprogramming of mitochondrial function and oxidative stress resilience (OxSR) through a process described as “mitohormesis” (*46–48*). In biological systems, hormesis describes a biphasic dose response in which a stress/signal is moderated by a compensatory response (*49*). For example, cancer cells produce more ROS than healthy cells, but the oncogene induced overproduction of ROS elicits compensatory ROS quenching OxSR response mediated by the transcriptional activity of NRF2 (nuclear factor erythroid 2–related factor 2). This NRF2-mediated ROS quenching transcriptional program facilitates metabolic remodeling that allows cancer cells to harness ROS-mediated proliferation and migration effects without succumbing to ROS-mediated cell death (*1*).

### Stiff ECM forces HSF1-mediated mitochondrial reprogramming

Since MECs cultured on stiff ECM experienced a greater amount of ROS stress we sought to determine if they had induced the NRF2-mediated Oxidative Stress Response (OSR) to survive and adapt. To test this, we utilized RNA-Seq to characterize the transcriptional state of MECs cultured on soft-to-stiff ECM or soft ECM with hyperglycemia for 24 hours. We compared the transcriptional programs induced by hyperglycemia and adhesion substrate elasticity because they both affect cytoskeletal tension, mitochondrial reorganization (fragment/toroid), and mitochondrial oxygen consumption. Unsupervised hierarchical clustering demonstrated that the greatest transcriptomic signature overlap occurred in MECs plated on the soft ECM (400 Pa) that were exposed to hyperglycemia [25 mM] and MECs plated on the stiff ECM (60k Pa) with physiological glucose [5 mM] (Figure 4B). Gene ontology analysis indicated that the categories that were downregulated in response to stiff ECM included oxidation-reduction process, oxidoreductase activity, ion transport, mitochondrion, and mitochondrial inner membrane (Figure S4A). We also found that of all mitocarta 2.0 (*50*) annotated mitochondrial genes encoded by the nuclear genome which change (up or down) in response to hyperglycemia or stiff ECM, the vast majority of these changes were conserved between both stress (Figure 4C). Specifically, a number of ETC subunits were downregulated (*NDUFA7, ATP5B, ATP5D, COX6b1* etc.) while mitochondrial-localized chaperones and proteases, which facilitate mitochondrial import, protein folding, and structural remodeling of the mitochondria during UPR^mt^ were upregulated (*YME1L1, HSPE1, DNAJC10 (hsp40)), HSPD1, HSPA9, HSPB11*, etc.) (*51*). With regards to NRF2, unexpectedly, cells cultured on stiff ECM had downregulated canonical ARE-containing target genes (HMOX1, TXN, GPX2, GPX4, and NQ01, etc).

**Figure 4:**
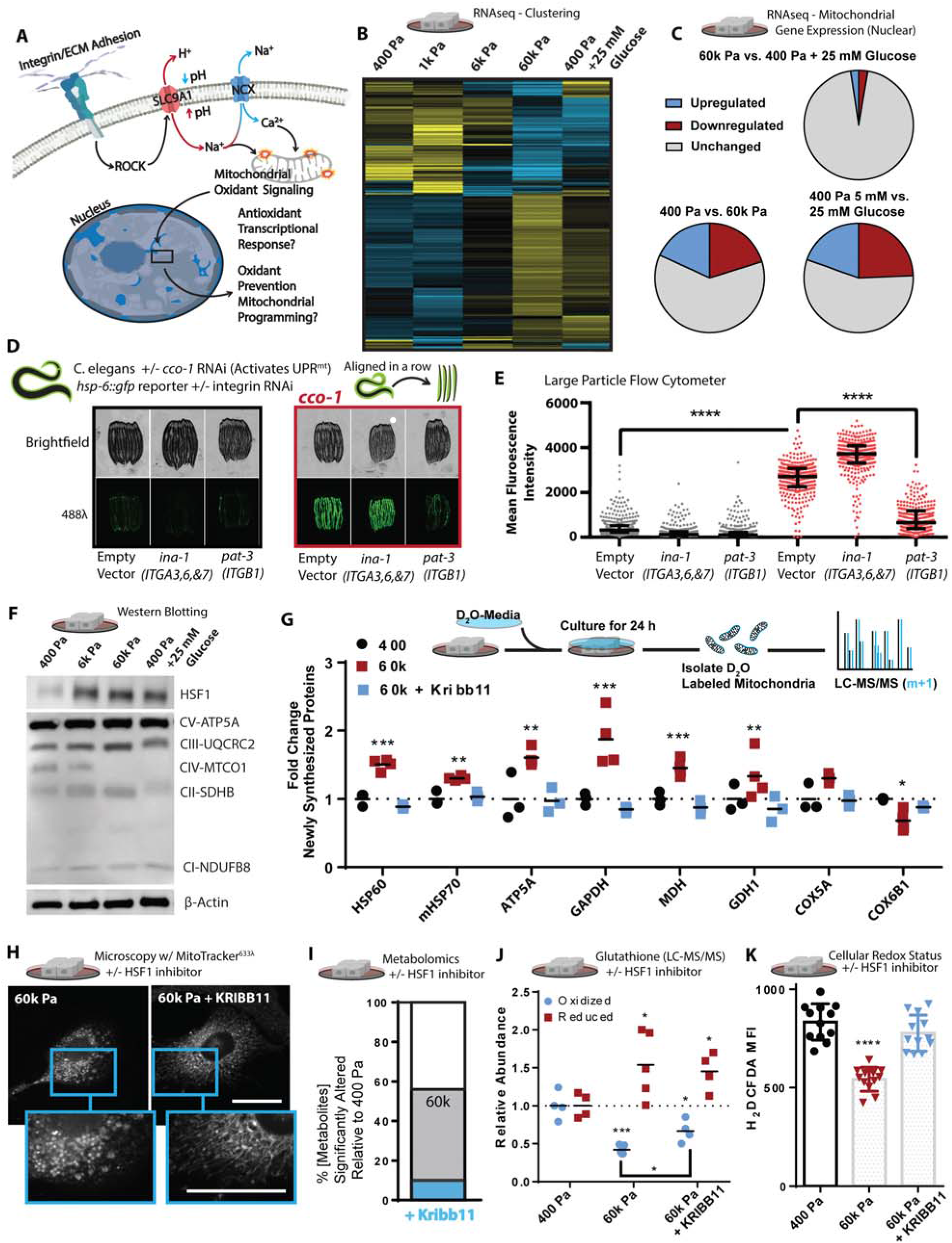
Mechanosignaling facilitates mitochondrial stress response *via* HSF1. A. Graphical representation of the paradigm and remaining questions. B. Heatmap depicting unsupervised hierarchical clustering of RNAseq of cells cultured on soft-to-stiff ECM for 24 h +/- glucose [5 or 25 mM], (n=2 duplicate libraries of 3 biological replicates, ~ 10 million reads per library). C. Comparison of significantly altered MitoCarta 2.0 catalogued genes from the 400 pa, 60k, and 400 pa + 25 mM glucose conditions shown in Figure 4B. D. *hsp-6::gfp* reporter fluorescent intensity representative images of *C. elegans* quantified, in Figure 4E RNAis were mixed at a 5:1 ratio of ev, *ina-1*, or *pat-3* RNAi to ev or *cco-1* RNAi as depicted. E. Quantification of *hsp-6::gfp* reporter fluorescent intensity of *C.elegans* measured with a large particle cytometer, +/- *cco-1* RNAi (n=387, 309, 377, 326, 312, and 294 animals in order left to right, repeated 3 times). F. Western blot depicting relative protein abundance of HSF1, electron transport chain components, or β-actin within 5 μg of total protein derived from lysates of cells cultured on soft-to-stiff ECM for 24 h +/- glucose [5 or 25 mM]. G. Stable Isotope mitochondrial proteomics of crude mitochondrial fraction of MCF10A cells grown 400 or 60k Pa ECM for 24 h +/- KRIBB11 [2 μM] [(n=4 biological replicates LC-MS/MS run together, repeated 2 times) H. Confocal microscopy of 100 nM mitotracker (deep red FM) stained and fixed (PFA) cells cultured on 60k Pa ECM surfaces for 24 h +/- vehicle or KRIBB11 [2 μM]. I. Metabolomics (LC-MS/MS) of cells cultured on 400 or 60k Pa ECM for 24 h, % significantly altered relative to 400 Pa +/- KRIBB11 [2 μM]. (n=4-5 biological replicates LC-MS/MS run together, repeated 2 times) J. Oxidized/reduced glutathione (NEM protected) measurements of MCF10A cells grown on 400 or 60k Pa ECM for 24 h +/- KRIBB11 [2 μM] [(n=4 biological replicates, repeated 2 times) (n=4-5 biological replicates LC-MS/MS run together, repeated 2 times). K. Oxidative stress indicator intensity of cells after 1 hour, MCF10A cells cultured on varied 400 or 60k Pa ECM for 24 h +/- vehicle or KRIBB11 [2 μM] prior, measured with 2’,7’-dichlorodihydrofluorescein diacetate (H_2_DCFDA) [2 μM]. *Data shown represents ± SEM. *P < 0.05, **P < 0.01, ***P < 0.005, < 0.0001 via oneway ANOVA with Tukey test for multiple comparisons in (E, G, J and K*).

It was paradoxical that genes with AREs in their promoters were not upregulated in MECs cultured on stiff ECM, since they had experienced a greater amount of redox stress (Figure 3E). However, this result could indicate that the transcriptomes measured reflected a post OxSR state, in which oxidants were not actively produced and therefore ARE-mediated gene expression was downregulated. We then compared the transcriptional signature of MECs cultured on stiff ECM to other known stress responses (*52*) that can remediate damage resulting from oxidative stress, such as protein misfolding (*53*). We found that the majority of genes which characterize the Integrated Stress Response (ISR) were downregulated, Heat Shock Response (HSR) genes were upregulated, and genes ascribed to the OSR and UPR^mt^ were inconsistently up- and downregulated. However, when comparing the upregulated genes of the UPR^mt^, OXR, and HSR, we noted that the upregulated genes were primarily heat shock proteins (HSPs) regulated by heat shock factor 1 (HSF1) (Table S1) (*52*).

Since activation of UPR^mt^ occurs in response to mitochondrial respiratory dysfunction (Figure 1B, 1E, and 1F), we tested if integrin signaling affected the activation of the UPR_mt_. We used cytochrome c oxidase-1 subunit Vb (*cco-1/COX4*) knockdown, which robustly induces a fluorescent UPR^mt^ reporter, *hsp-6::gfp (HSPA9* orthologue), in *C. elegans*. We performed double RNAi of *cco-1* in conjunction with knockdown of *ina-1* (ITGA orthologue, most similar to *ITGA3,6, and 7*) or *pat-3* (*ITGB1* orthologue). We found that *pat-3/ITGB1* knockdown robustly attenuated *cco1*-mediated UPR^mt^, which suggests that integrin signaling is an important input to the activation of UPR^mt^ (Fig. 4D-E). We then determined that HSF1 is required for the maximal activation of *cco1*-mediated UPR^mt^ in *C.elegans*, as RNAi-knockdown of *hsf-1* partially suppressed UPR^mt^ induction (Figure S4B). Since *C. elegans* activate UPR^mt^ primarily through *atfs-1* (*54*), which does not have a clear mammalian orthologue (*55*), the data suggest HSF1 may play a larger role in the mammalian UPR^mt^ (*56*) or could resolve a UPR^mt^-overlapping aspect of mitochondrial dysfunction (*57*).

HSF1 is primarily known to regulate a transcription program that facilitates the survival of cells experiencing heat stress (~43°C), but it may have an unappreciated role in OxSR since its transcriptional activity is regulated by ROS (H_2_O_2_) (*58*). Upregulation of HSF1 expression is an outcome of the NRF2-mediated OSR, because the HSF1 promoter (−1.5k to −1.7k bp) is heavily enriched with AREs (*59*). Indeed, we found that HSF1 abundance increased in response to stiff ECM or hyperglycemia, as was mitochondrial ATP5A, but not the mitochondrial encoded subunit of oxygen consuming ETC complex IV subunit MTC01 (Figure 4F). Inhibiting SLC9A1 in MECs cultured on stiff ECM repressed the expression of HSF1 and its downstream targets (Figure S4C). Treatment with MitoTEMPO, a mitochondria-targeted antioxidant, suppressed stiff ECM or paraquat-induced HSF1 and HSF1-target gene expression (Figure S4D and E). Overall, this indicates that mitochondrial ROS induced by the stiff ECM *via* SLC9A1 (Figure 3E and S3I) activity promotes HSF1 expression and activity.

HSF1 could modify mitochondrial structure/function by influencing the expression of the mitochondrial import machinery, such as mtHSP70 (*HSPA9*) (*60*), or by regulating mitochondrial biogenesis in collaboration with peroxisome proliferator-activated receptor gamma coactivator 1-alpha (PGC-1α) (*61*). To test this possibility, we measured the incorporation of newly synthesized proteins into the mitochondria with stable isotope incorporation mass spectrometry, which revealed that the stiff ECM enhanced the incorporation of newly synthesized proteins in an HSF1-dependent fashion (Figure 4G). Consistent with the hypothesis that an HSF1-mediated response facilitated the stiff ECM induced mitochondrial adaptation, HSF1 inhibition was sufficient to prevent the altered mitochondrial morphology induced by stiff ECM (Figure 4H). Inhibition of HSF1 also restored ~80% of the metabolite concentrations measured in MECs plated on stiff ECM to that of MECs cultured on soft ECM (Figure 4I and S4F). Overall, these findings suggest that ECM mechanosignaling alters mitochondrial reorganization and metabolic programming through a heat-stress independent HSF1-mediated program (*62*).

To explore if adhesion-mediated mechanosignaling facilitates a HSF1-dependent OxSR program, we quantified the levels of reduced and oxidized glutathione, the primary cellular oxidant detoxification and redox (reduction:oxidation) management system which becomes oxidized in the presence of ROS. MECs cultured on stiff ECM had lower levels of oxidized glutathione than those on soft ECM. HSF1 inhibition was sufficient to significantly increase the levels of oxidized glutathione in MECs on stiff ECM (Figure 4J). Metabolomics allowed us to observe that many metabolite changes that reflect OxSR (*63*), such as pentose phosphate pathway activity, which generates reduced nicotinamide adenine dinucleotide phosphate (NADPH) required to regenerate reduced glutathione and mitigate oxidative stress, were also elevated in response to stiff ECM, and could be normalized by inhibiting HSF1 (Figure S4F). Indeed, HSF1 inhibition abolished the enhanced reducing capacity of MECs that had been cultured on stiff ECM for 24 hours prior to the measurement of redox stress (Figure 4K). These data indicate that while MECs experience more redox stress in response to stiff-ECM, they adapt and acquire OxSR through HSF1-dependent changes in cellular metabolism facilitated by mitochondrial reprogramming *via* compositional changes (Figure 1D, 4E-F, and 5A).

**Figure 5:**
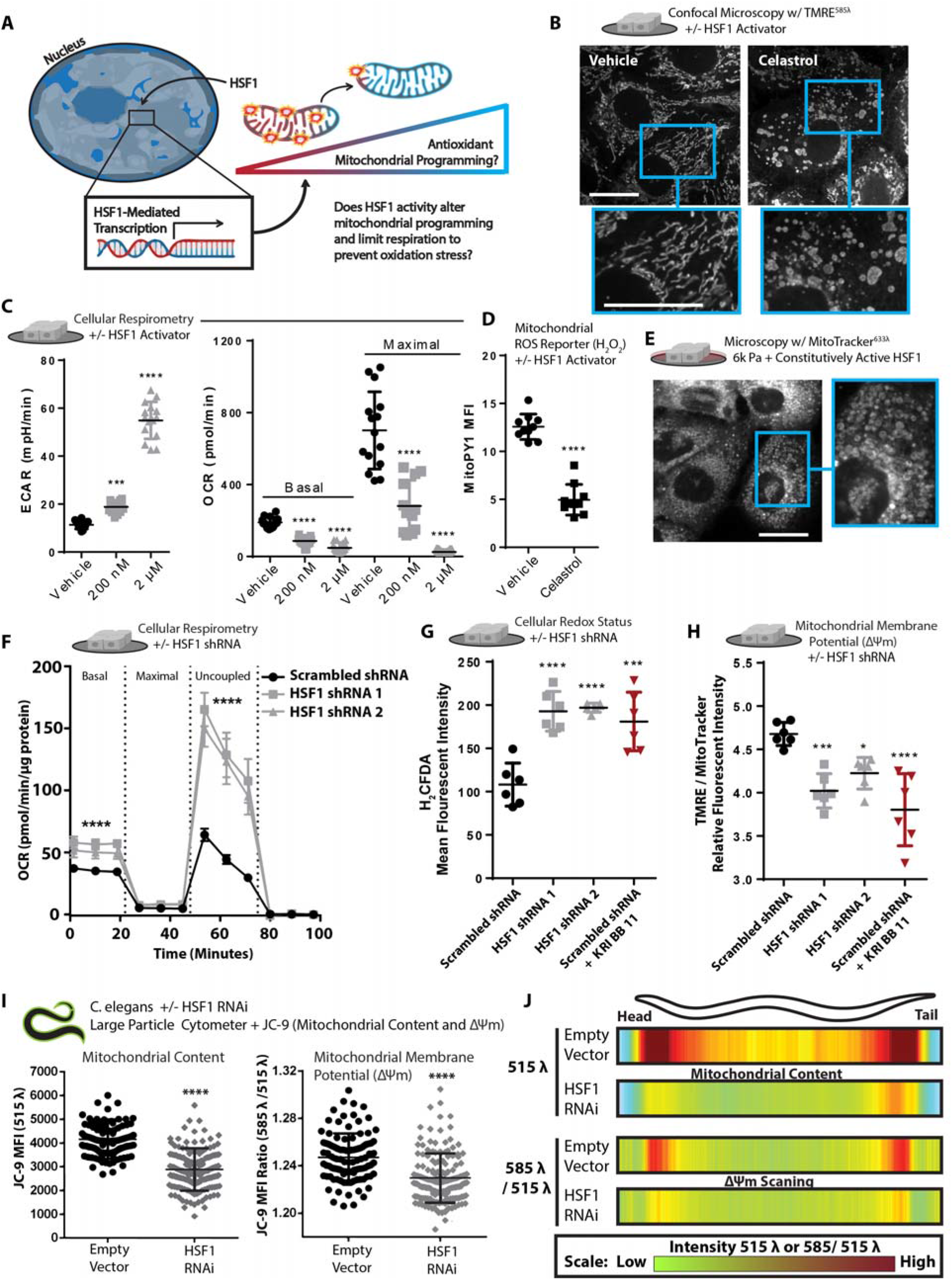
HSF1 induces mitochondrial reprogramming. A. Graphical depiction of experimental question. B. Confocal microscopy depicting morphology and mitochondrial membrane potential staining of live cells *via* TMRE [10 nM] staining +/- vehicle or Celastrol [2 μM] treatment for 40 minutes prior to imaging. C. Extracellular acidification rate (ECAR) and OCR of MECs treated +/- vehicle or Celastrol [200 nM] for 24 h or Celastrol [2 μM] for 40 minutes (n=5 wells, 3 replicate measures). D. B. Mitochondrial H_2_O_2_ production of cells treated with Celastrol [2 μM] treatment for 40 minutes, measured with MitoPY [1 μM] (n=5, repeated 2 times). E. Confocal microscopy depicting mitochondrial morphology of PFA-fixed cells expressing constitutively-active HSF1 and cultured on 6k Pa ECM for 24 h, stained with 100 nM mitotracker (deep red FM). F. Oxygen consumption rate (OCR) of MCF10A cells expressing a scrambled shRNA or two different shRNAs targeting HSF1 (n=5 wells, 3 replicate measures, repeated 3 times) G. Oxidative stress indicator intensity after 1 h in cells expressing a scrambled shRNA +/- KRIBB11 [2 μM] or two different shRNAs targeting HSF1, measured with 2’,7’- dichlorodihydrofluorescein diacetate (H2DCFDA) [2 μM]. (n=6 wells, repeated 3 times). H. Mitochondrial membrane potential of MCF10A cells expressing a scrambled shRNA +/- KRIBB11 [2 μM] or two different shRNAs targeting HSF1, measured with TMRE [1 nM] and mitotracker [100 nM] after 1 h staining (n=6, repeated 3 times). I. Mean fluorescent intensity of 150 per condition JC-9 stained *C. elegans* grown on empty vector or *hsf-1* RNAi from hatch, depicting mitochondrial mass (515 λ alone) or mitochondrial membrane potential (585 λ / 515 λ), spatially quantified in Figure 5J. J. Heatmap depicting mitochondrial mass (515 λ alone) or mitochondrial membrane potential (585 λ / 515 λ) across the body length (head (left) to tail (right)) of 150 *C. elegans* animals grown on empty vector or *hsf-1* RNAi from hatch; JC-9 staining *via* administration of JC-9 loaded *C.elegans* food (E.coli) (repeated 3 times). *Data shown represents ± SEM. *P < 0.05, **P < 0.01, ***P < 0.005, < 0.0001 via twotailed unpaired Student t test in (D, F, and I) and one-way ANOVA with Tukey test for multiple comparisons in (C, G, and H*).

HSF1 can be pharmacologically activated using Celastrol, a reactive electrophile derived from the “Thunder of God” vine (Tripterygium wilfordii) (*64*), which was sufficient to induce mitochondrial fragmentation/toroids in MECs on all substrates (Figure 5B). HSF1 activation also increased extracellular acidification (ECAR), a proxy measure of glycolytic flux, and reduced mitochondrial oxygen consumption (Figure 5C). A mitochondria-localized ROS (H_2_O_2_) reporter (MitoPY1) revealed that MECs treated with Celastrol had significantly suppressed mitochondrial ROS production (Figure 5D). Expression of a constitutively active HSF1 induced fragmented/toroidal mitochondria in MECs plated on the soft ECM (Figure 5E and S5B), and increased mitochondrial membrane potential (Figure S5C). Conversely, HSF1 knockdown increased mitochondrial respiration (Figure 5F), induced oxidative stress (Figure 5G & S5D), and decreased mitochondrial membrane potential (Figure 5H & S5E) in both the MCF10A (nonmalignant) and MDA-MB-231 (aggressive and malignant) MECs (Figure S5F-I). The physiological relevance of these findings was confirmed by finding that reducing *hsf-1* expression in *C. elegans* decreased mitochondrial content and membrane potential throughout the whole organism (Figure 5I-J). This indicates that increased HSF1 activity reduces mitochondrial oxygen consumption and increases the mitochondrial membrane potential because proton flux from IMS to MM is reduced, which may limit ROS produced as a byproduct of OxPhos to facilitate enhanced OxSR.

### ECM-mechanosignaling engenders mitochondrial OxSR *via* HSF1 and YME1L1

Thus far, our data suggested that adhesion-mediated mechanosignaling stimulates a heat-stress independent HSF1 transcriptional program, previously implicated in cancer, that alters mitochondrial structure/function, restricts mitochondrial respiration, and oxidant production. Accordingly, we next stress-tested if the OxSR adaption was sufficient to oppose mitochondrial ROS-mediated apoptosis, a trait associated with many tumors (*65*). We treated MECs cultured on soft-to-stiff ECM, with paraquat and assayed for apoptosis (*31*). Consistent with the hypothesis that mechnosignaling promotes OxSR *via* HSF1, MECs cultured on stiff ECM were less sensitive to paraquat treatment. This OxSR phenotype could be further enhanced *via* expression of constitutively active HSF1 and ablated by HSF1 knockdown or inhibition (Figure 6A-B & S6A-B). Functional links between mechanosignaling and mitochondrial OxSR adaptation were verified by determining that mitochondrial depletion negated the impact of ECM stiffness on redox sensitivity to paraquat (Figure S6C).

**Figure 6:**
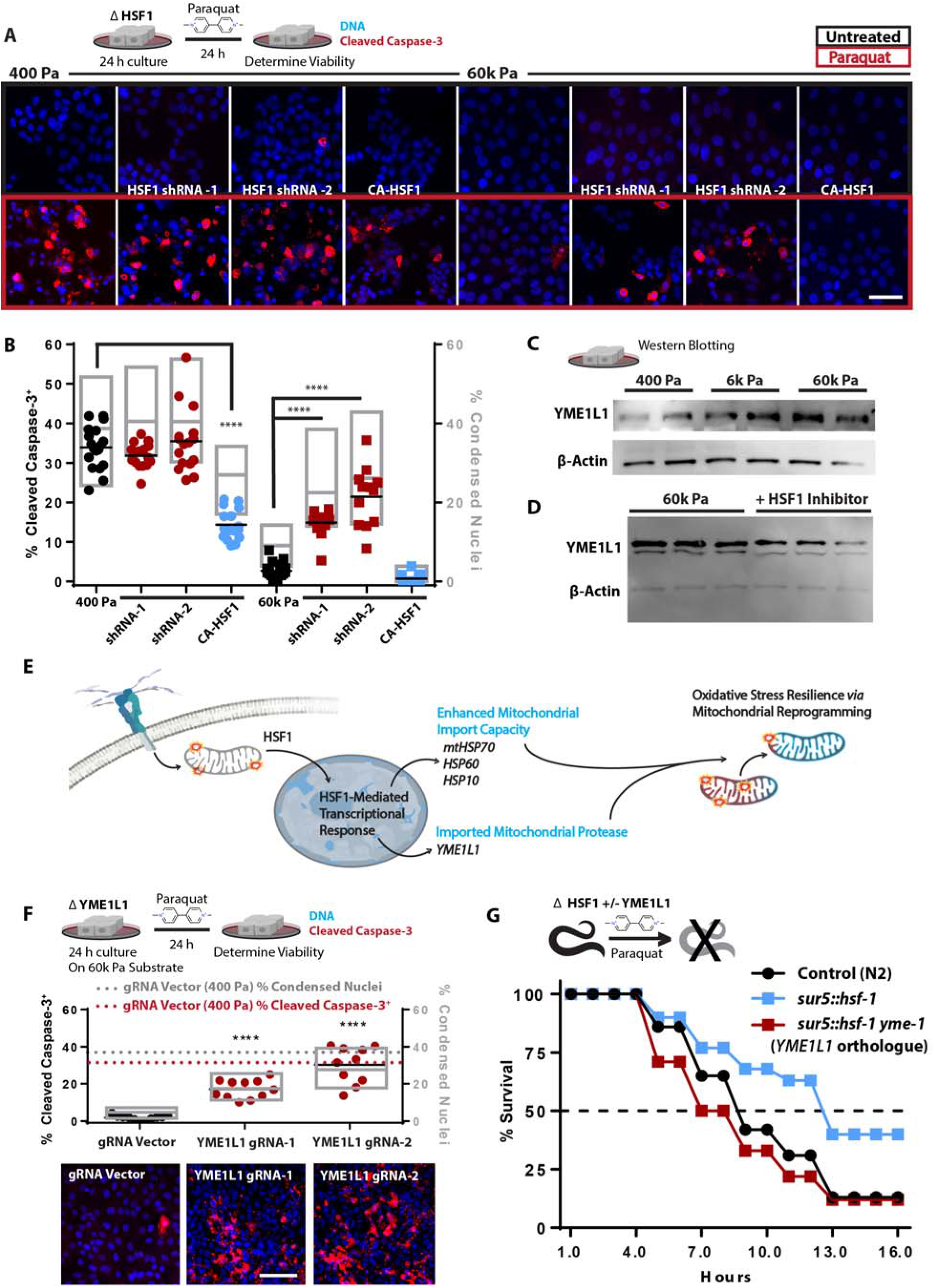
ECM-mediated mechanosignaling controls OxSR *via* HSF1 and YME1L1. A. Confocal microscopy of indicators of apoptosis with cleaved caspase 3 staining (red) and nuclear condensation (dapi) of MECs cultured on 400 or 60k Pa ECM for 24 h with subsequent 24 h +/- paraquat treatment [10 mM]. 100k cells/well of 24 well plate. (n=4 replicates, repeated 3 separate times) B. Quantitation of cells from 16 field views depicted in G for condensed nuclei and cleaved caspase-3 positive cells (1653-575 cells counted per condition, repeated 3 times). C. Western blot of YME1L1 and β-actin from 5 μg of protein derived from cells cultured on soft-to-stiff ECM for 24 h. (2 biological replicates shown, repeated 3 times) D. Western blot of YME1L1 and β-actin from 5 μg of protein derived from cells cultured on 60k Pa ECM for 24 h +/- KRIBB11 [2 μM] (3 biological replicates shown, repeated 2 times) E. Graphical representation of the conceptual paradigm pertaining to this figure. F. Quantitation of MECs with YME1L1 knockdown *via* CRISPR-I compared to CRISPR-I and empty guide vector expressing cells on 400 Pa (dashed lines) or 60k Pa ECM, 11 field views quantified for condensed nuclei and cleaved caspase-3 positive cells (923-2880 cells counted per condition, repeated 3 times). G. *C. elegans* survival in 50 mM paraquat, with *C. elegans* overexpressing *hsf-1 (sur-5p::hsf-1*) compared or control line (N2) grown on either empty vector or *ymel-1* RNAi from hatch (n=80 animals per condition, repeated 3 times). *Data shown represents ± SEM. *P < 0.05, **P < 0.01, ***P < 0.005, < 0.0001 via oneway ANOVA with Tukey test for multiple comparisons in (B and F*).

Due to the fact that stiff ECM and HSF1 altered mitochondrial protein turnover rates, and mitochondrial import is required for mitochondrial protein turnover, we decided to test if cytoskeletal tension mediated OxSR was dependent upon mitochondrial protein import. Nuclear-encoded mitochondrial proteins are imported into the mitochondria (*66*), a process which requires a number HSF1 target genes to efficiently occur (e.g. *HSPD1* (HSP60), *HSPE1* (HSP10), and *HSPA9* (mtHSP70) (*67, 68*). To test if cytoskeletal tension enhanced mitochondrial import was critical for OxSR, we used JG-98, an inhibitor of HSP70 that enriches in mitochondria (*69, 70*). JG-98 treated cells grown on stiff ECM were not apoptotic, but were severely sensitized to paraquat-induced death (Figure S6D). To identify the specific mediators of the HSF1-dependent OxSR we cross-referenced conserved nuclear-encoded mitochondrial genes containing heat-shock elements (HSE) in their promoters with genes whose expression was upregulated by cytoskeletal tension (Figure 3A). We found one likely candidate, mitochondrial escape 1 like 1 (*YME1L1*), a zinc-dependent metalloprotease of the AAA^+^ protein family (ATPases with diverse cellular activity), which is a hallmark of UPR^mt^ and regulates mitochondrial morphology (*71, 72*). We verified that YME1L1 protein levels were regulated by HSF1 and were upregulated in response to mechanosignaling *via* stiff ECM (Figure 6C-D).

Since mitochondrial import stress activates HSF1 (*57*), and UPR^mt^ upregulates the import machinery, we hypothesized that dysfunctional or ROS-producing mitochondria promote mitochondrial import capacity to facilitate the import of certain nuclear-encoded mitochondrial proteins that mediate mitochondrial repair/reprogramming (Figure 6E). To determine if YME1L1 played a role in stiff ECM induced OxSR, we utilized CRISPR-I to downregulate YME1L1 expression. YME1L1 knockdown sensitized MECs cultured on stiff ECM to paraquat-induced death almost as much as those cells plated on soft ECM (Figure 6F). To verify that HSF1 activity conferred OxSR through YME1L1, we examined the paraquat sensitivity of *hsf-1-* overexpressing C. elegans with or without reduced expression of *yme-1 (YME1L1* orthologue). Overexpression of *hsf-1* rendered *C. elegans* more resistant to paraquat-induced death, to a greater extent than that observed in other long-lived strains (e.g. daf-2 knockdown). Impressively, *yme-1* knockdown completely abolished the OxSR conferred to C. elegans through the overexpression of *hsf-1* (Figure 6G). Overall, the data indicate that YME1L1 plays an essential role in the HSF1-mediated OxSR that is induced by stiff ECM induced adhesion-mediated mechanosignaling.

## Discussion

We identified a mechanism whereby the physical properties of the microenvironment alter mitochondrial composition, structure and function to tune cellular metabolism through a stress adaptation. We demonstrate that SLC9A1 and HSF1 alter mitochondrial function and support OxSR by regulating the levels of YME1L1 (*72*). The mechano-responsiveness of HSF1 and its ability to limit mitochondrial respiration, may explain why oncogene driven Warburg metabolism has been so difficult to observe *in vitro*. The rigid tissue culture polystyrene substrates (3G Pa) elevates mechanosignaling and chronically activates HSF1, regardless of oncogene-transformation, and this effect obscures any comparative measurements of mitochondrial function in normal and oncogene-transformed cells. Instead, prudent use of model systems with biomimetic properties (physically and chemically similar to the relevant biological system) are needed to uncover oncogene driven alterations in mitochondrial metabolism (*73, 74*).

Our findings demonstrate that HSF1-driven redox management not only suppresses the production of ROS, by limiting mitochondrial respiration, but it also opposes oxidant damage by promoting mitochondrial biogenesis/protein turnover and enhancing reducing equivalents (glutathione/NADPH). Previous studies have indicated that cell-detachment/attachment-associated signaling elicits redox stress (*75–77*). With that in mind, coupling redox stress management to a molecular rheostat would be a rational design principle. HSF1 is a logical candidate to serve as such a molecular rheostat. If HSF1 is such a molecular rheostat it would become transcriptionally active when misfolded proteins accumulate. Misfolded proteins can accumulate due to changes in pH, ion concentrations, osmolality, osmotic pressure, molecular crowding, adhesion-associated forces (mechanotransduction), enthalpy (heat), entropy (order), and redox balance (*58, 78–81*) - all of which are cellular conditions associated with HSF1 activation. By surveying the physical state of the proteome, HSF1 is well poised to temper diverse environmental perturbations that elicit mitochondrial dysfunction and oxidant leak. Indeed, HSF1 could mitigate the redox stress induced by conditions that deform mitochondrial structure (*82*), such as the physical stresses cells encounter in tumors with high interstitial pressure, mechanically stressful metastatic sites (*83*), rigid ECMs, or oncogene induced ROCK activity (*84–86*).

HSF1 levels are elevated in the majority of tumors and is implicated in cancer aggression and metastasis (*62, 87, 88*). Because tumors are stiffer than healthy adjacent tissues, our findings offer a tractable explanation for why HSF1 and its target genes are so frequently upregulated in tumors (*89*). The heat shock independent activation of HSF1 and its target genes would provide the tumor cells with a metabolic adaptation to this chronic mechanical stress (Figure S6E-F). In this regard, therapeutic approaches to disrupt HSF1 and its target genes have focused on cytosolic and nuclear targets, but can incur difficult to tolerate systemic effects in humans (*90*). Our data implicate the mitochondrial protease YME1L1 as a HSF1 target gene that facilitates mitochondrial redox stress adaptations, which could be pharmacologically targeted as an anti-tumor therapeutic (*72*).

## Acknowledgments

We thank the following people for their support and advice: Diane Barber and Yi Liu, provided advice and reagents pertaining to SLC9A1 and pH regulation. Milos Simic, generated the constitutively active HSF1. Gilberto Garcia, developed the MLS mRuby C.elegans strain. Todd McDevitt, Serah Kang, and Ariel Kauss provided editorial suggestions which significantly improved the quality of the manuscript. Chris Phillart, maintained the cellular respirometer used in these studies. Brant Webster, generated the MTS-roGFP2 vector. Fui Boon Kai, acquired the ROCK:ER construct and generated the ROCK:ER stable cell line. Bram Piersma, provided dry Dutch humor, beneficial laboratory ambiance, and conversation critical to the development of this manuscript.

## Author contributions

Conceptualization: K.M.T. and V.M.W. Methodology: K.M.T, R.H.S. G.T., C.G.C., B.F., V.M.W., & A.D. Investigation: K.M.T., R.H.S., G.T., C.G.C., B.F., C.S., J.M., C.S., J.R.D., S.S.M., K.C. Formal Analysis: K.M.T., R.H.S., G.T., B.F.,A.L.R., D.L.S., C.S., J.M., C.S., J.R.D., S.S.M. Data curation: K.M.T. Funding acquisition: V.M.W., K.M.T, A.D., M.H., D.K.N., A.R.D., K.S., J.G, D.L.S, & N.J.K. Project administration: K.M.T. Resources: V.M.W., K.M.T, A.D., M.H., D.K.N., A.R.D., K.S., H.S., & J.G. Software: C.S. & P.F. Supervision: V.M.W., A.D., K.S., A.R.D., D.K.N., D.LS., M.H., J.G. Validation: K.M.T., R.H.S., G.T.,A.S.M., C.G.C, and C.S. Visualization: K.M.T. Writing – original draft: K.M.T. Writing – review & editing: V.M.W., K.M.T., A.R.D., J.R.D., R.H.S., G.T., A.S.M, C.G.C., J.M., C.S., S.S.M.

## Materials and Methods

### Cell culture

MCF10A and MB-MDA-231 cells were sourced from ATCC, routinely tested and found to be free of mycoplasma contamination, and maintained below passage 22. All cells were maintained and in 5% CO^2^ at 37 °C. MCF10A were cultured in [5 mM] glucose DMEM:F12 (1:1 mixture of F:12 [10mM] glucose and [0 mM] glucose DMEM) (Life Technologies, 11765054 and 11966025) supplemented with 5% Horse Serum (Gibco, 16050-122), 20 ng/mL epidermal growth factor (Peprotech), 10 μg/mL insulin (Sigma), 0.5 μg/mL hydrocortisone (Sigma), 100 ng/mL cholera toxin (Sigma, C8052-2MG), and 1x penicillin/streptomycin (Gibco). MB-MDA-231 tumor cells (ATCC) were grown in 5 mM glucose DMEM supplemented with 10% fetal bovine serum (FBS) (Hyclone) and 1x penicillin/streptomycin. HEK293T cells (ATCC) were maintained in DMEM supplemented with 10% FBS and 1x penicillin/streptomycin and were used to produce lentiviral particles with psPAX2 (Addgene 12260), pMD2.G (Addgene 12259) and various transfer vectors described hereafter (https://www.addgene.org/guides/lentivirus/).

### ECM coated polyacrylamide hydrogel cell culture surfaces (PA-gels)

Cleaned (10% ClO, 1M HCL, then 100% EtOH) round #1 German glass coverslips (Electron Microscopy Services) were coated with 0.5% v/v (3-Aminopropyl)triethoxysilane (APTES, Sigma, 440140), 99.2% v/v ethanol, and 0.3% v/v glacial acetic acid for 2 h and then cleaned in 100% EtOH on an orbital shaker at 22 °C. APTES activated coverslips were coated with PBS buffered acrylamide / bis-acrylamide (Bio-Rad, 1610140 and 1610142) solutions (3% / 0.05% for 400 Pa, 7.5% / 0.07% for 6k Pa, and 10% / 0.5% for 60k Pa) polymerized with TEMED (0.1% v/v) (Bio-Rad, 1610801) and Potassium Persulfate (0.1% w/v) (Fisher, BP180) to yield a final thickness of ~ 85 μm. PA-gels were washed with 70% EtOH and sterile PBS prior 3,4-dihydroxy-L-phenylalanine (DOPA) coating for 5 min at 22 °C protected from light with sterile filtered DOPA in pH 10 [10 mM] Tris buffer (*91*). DOPA coated PA-gels were washed 2x with sterile PBS and ECM functionalized with 5 μg/mL human fibronectin (Millipore, FC010) in sterile PBS 1 h at 37 °C to generate an expected fibronectin coating density of 6 μM/cm^2^.

### Immunofluorescence microscopy

Cells or tissues were fixed in 4% paraformaldehyde (Electron Microscopy Services, 15710) in 1X PBS for 30 min at room temperature, washed and blocked with a blocking buffer (HBSS fortified with: 10% FBS (Hyclone), 0.1% BSA (Fischer, BP1600), 0.05% saponin (EMD, L3771), and 0.1% Tween 20 (Fischer, BP337500). Primary antibodies [1:100-1:200] for 2 h at RT or 24 h at 4 °C, Secondary antibodies [1:1000] for 2 h at 22 °C. Samples were imaged with a Nikon Eclipse Ti spinning disc microscope, Yokogawa CSU-X, Andor Zyla sCMOS, Andor Multi-Port Laser unit, and Molecular Devices MetaMorph imaging suite.

Antibodies used: Cleaved Caspase-3 (Asp175) (Cell Signaling, 9661, AB_2341188), HSF1 (Cell Signaling, 4356, AB_2120258), and HSP60 (LK1, sc-59567, AB_783870).

### MitoTacker staining

Mitotracker deep red FM or Mitotracker Green FM (Invitrogen, M22426 and M7514) was solubilized in DMSO to yield 100 μM frozen aliquots which were diluted into media to yield a 10 μM stock which was added directly to cell culture media already in the culture yielding a final concertation of 100 nM (to prevent media change derived fluid flow shear stress) 30 min before 4% PFA fixation or live cell imaging (MitoTracker red FM was used for PFA fixed samples).

### TMRE, H2DCFDA, Rhod-2 AM, Calcium Green-1 AM, and MitoPy1

Stained cells were washed twice with PBS and imaged on a SpectraMax i5 Multi-Mode plate reader. Experiments were carried out with 50k or 100k cells per well in 500 μL of media in 24 well format with or without fibronectin coated PA-gels (indicated). 2 μM Rhod-2 AM (Thermo, R1244) and 2 μM Calcium green-1 AM (Life Technologies, C3011MP) was added to culture media and allowed to stain at 22 °C for 20 min prior to imaging (frozen Rhod-2 AM aliquots were used only when the DMSO suspension remained clear), or 2 μM H_2_DCFDA (Thermo, D399) was added to media 1 or 4 h prior to imaging, or 5 μM MitoPy1 (Tocris, 4428) was added to media 1 h prior to imaging, 2 nM TMRE (Fischer, T669) was applied to cells in media without disturbing the existing culture media (similar dilution scheme to MitoTracker) for 1 h prior to microscopy or plate reader assay.

### Intracellular pH (pH_i_)

10 μM BECEF (Invitrogen, B1150) was added to cell culture media for 30 min at 37 °C in 5 % CO_2_ incubator. Cultures were washed twice and then fluorescent intensities of BCECF was determined with a SpectraMax i5 plate reader in a buffer comprised of 25 mM HEPES, 140 mM NaCl, 5 mM KCl, 1 mM KHPO4, 1 mM MgSO4, 2 mM CaCl2, and 5 or 25 mM glucose. The cultures were then treated with a pH 7.7 buffer containing 10 μM Nigericin (Invitrogen, N1495), 25 mM HEPES, 105 mM KCl, and 1 mM MgCl for 5 min at 22 °C followed by determination of BCECF fluorescent intensities at high pH. The cultures were then treated with a pH 6.6 buffer containing 10 μM Nigericin (Invitrogen, N1495), 25 mM HEPES, 105 mM KCl, and 1 mM MgCl for 5 min at 22 °C followed by determination of BCECF fluorescent intensities at low pH. A linear relationship between pH 6.6 and 7.7 was observed and sample pH was estimated relative to pH standards for each individual culture well.

### qPCR

Total RNA was isolated from biological samples with TRIzol (Invitrogen, 15596-018) according to the manufacturer’s instructions. cDNA was synthesized with 1 μg total RNA in 10 μL reaction volume with RNA using M-MLV reverse transcriptase (BioChain, Z5040002-100K) and 5X reaction buffer (BioChain, Z5040002-100K), random hexamers (Roche, 11034731001), dNTPs, and 1U of Ribolock (ThermoFischer, EO0384). RT-thermocycler program: random hexamers and RNA incubated at 70°C for 10 min, then held at 4°C until the addition of the M-MLV reverse transcriptase, dNTPs, Ribolock, and M-MLV-reverse transcriptase, then 50°C for 1 h, 95 °C for 5 min, then stored at −20 °C until qPCR was performed. The reverse transcription reaction was then diluted to 50 μL total volume with ddH_2_O rendering a concentration of 20 ng RNA per 1 μL used in subsequent qPCR reactions. qPCR was performed in triplicate using PerfeCTa SYBR Green FastMix (Quantabio, Cat# 95072-05K) with an Eppendorf Mastercycler RealPlex^2^. qPCR thermocycler program: 95° C for 10 min, then 40 cycles of a 95°C for 15 s, 60°C for 20 s, followed by a melt curve 60-95°C over 10 min. Melt curves and gel electrophoresis were used to validate the quality of amplified products. The ΔCt values from independent experiments were used to calculate fold change of expression using the 2^-ŬΔCt^ method. For each gene measured, the SEM of the ΔCt values was calculated and used to generate positive and negative error values in the 2^-ΔΔCt^ fold change space. Plots of qPCR data display bars representing the mean fold change ±SEM and individual points representing the fold change value for each experiment relative to the mean.

### qPCR primers used

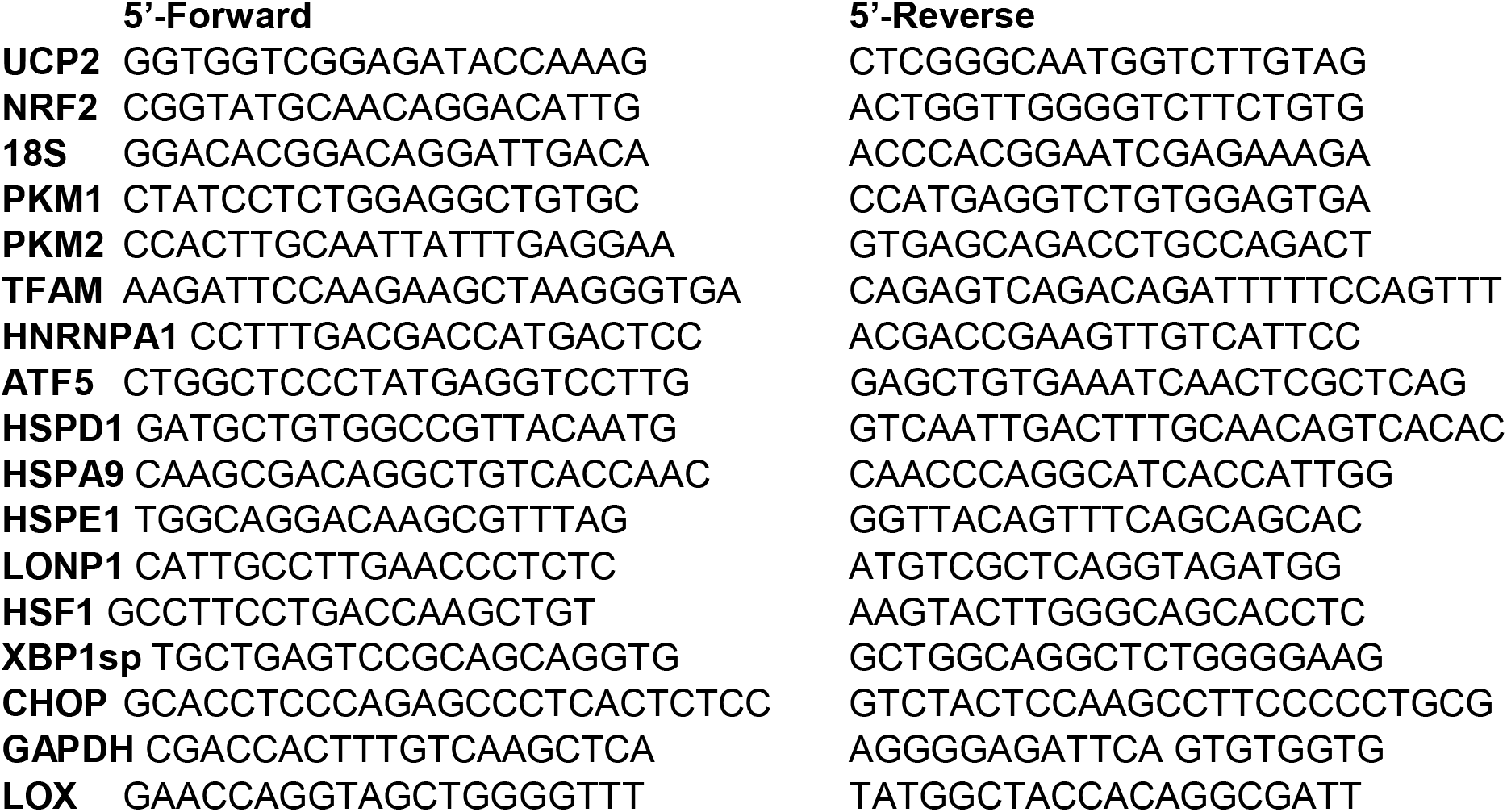

### Western blotting

Cells were freeze (−80 °C) thaw lysed with RIPA buffer (150 mM NaCl, 1% v/v NP-40, 0.5% w/v sodium deoxycholate, 0.1% w/v SDS, and 25 mM tris) containing protease and phosphatase inhibitor coctail (GenDepot, P3100 and P3200). Protein content was determined via BCA (Pierce, 23225) and 5-10 μg of protein was mixed with 5x Laemmli buffer to generate final 1x concentration (50 mM Tris-HCl (Fischer, AAJ2267636) pH 6.8, 4% w/v SDS (Sigma, L3771), 10% v/v glycerol (Fischer, BP229-1), 0.1% w/v bromophenol blue (Bio-Rad, 1610404), 2% v/v β-mercaptoethanol (Bio-Rad, 1610710) and heated to 95 °C for 5 min (no heating of the samples used for the total oxphos (abcam, ab110413) blots). 10%-gels (Bio-Rad, Bulletin_6201) were cast in a PROTEAN Plus multi casting chamer (Bio-Rad). Samples were loaded (~20 μL) and run to completion in Tris Glycine SDS running buffer (25 mM Tris, 192 mM glycine (Fischer, BP381), and 0.1% SDS, pH ~8.6), wet transferred @ 100V for 60 min to methanol (Fischer, A412) activated PVDF (BioRad, 1620177) in Towbin transfer buffer containing (25 mM Tris, 192 mM Glycine, 20% v/v methanol, pH ~8.3). Protein loaded PVDF membranes were washed 2x with TBST (20 mM tris, 150 mM NaCl (S271), 0.1% w/v Tween20) and blocked in 5% milk TBST buffer for 1 h at 22 °C on an orbital shaker.

Antibodies used: HSF1 (Cell Signaling, HSF1 4356), YME1L1 (Invitrogen, PA564299, HSP60 (LK1, sc-59567), HSP70 (3A3, sc32239), mtHSP70 ((D-9), sc-133137), Total OXPHOS WB Antibody Cocktail (ab110413), β-Actin (Sigma, A5441), FAK pY397 (Invitrogen, 44-625G), FAK (BD, 610088), and pMLCK (Cell Signaling, 3671)

### ROCK:ER

pBABEpuro3 ROCK:ER (*21*) was generously provided by Dr. Michael F. Olson, packaged into retroviral particles with phoenix cells, and used to generate stable MCF10A cells lines which were activated with 1 μg/mL 4-Hydroxytamoxifen (Sigma).

### MTS-roGFP2

*COX4L MTS* tagged roGFP was expressed under the CMV promoter in a puromycin lentiviral transfer vector generated by the Dillin Lab (Dr. Brant Webster).

ATGCTTGCCACTAGAGTCTTTTCATTGGTAGGTAAAAGGGCCATAAGTACATCAGTCTGCGTGAGAGCCCACACCGGACCGGTCAGCAAGGGCGAGGAGCTGTTCACCGGGGTGGTGCCCATCCTGGTCGAGCTGGACGGCGACGTAAACGGCCACAAGTTCAGCGTGTCCGGCGAGGGCGAGGGCGATGCCACCTACGGCAAGCTGACCCTGAAGTTCATCAGCACCACCGGCAAGCTGCCCGTGCCCTGGCCCACCCTCGTGACCACCCTGACCTACGGCGTGCAGTGCTTCAGCCGCTACCCCGACCACATGAAGCGGCACGACTTCTTCAAGTCCGCCATGCCCGAAGGCTACGTCCAGGAGCGCACCATCTTCTTCAAGGACGACGGCAACTACAAGACCCGCGCCGAGGTGAAGTTCGAGGGCGACACCCTGGTGAACCGCATCGAGCTGAAGGGCATCGACTTCAAGGAGGACGGCAACATCCTGGGGCACAAGCTGGAGTACAACTACAACTGCCACAACGTCTATATCATGGCCGACAAGCAGAAGAACGGCATCAAGGTGAACTTCAAGATCCGCCACAACATCGAGGACGGCAGCGTGCAGCTCGCCGACCACTACCAGCAGAACACCCCCATCGGCGACGGCCCCGTGCTGCTGCCCGACAACCACTACCTGAGCACCTGCTCCGCCCTGAGCAAAGACCCCAACGAGAAGCGCGATCACATGGTCCTGCTGGAGTTCGTGACCGCCGCCGGGATCACTCTCGGCATGGACGAGCTGTACAAGTAA

### V737N β1 integrin

Weaver lab generated (Dr. Jonathan Lakins) puromycin lentiviral transfer vector expressing 3x myc tagged V737N β1 integrin using the tetracycline rtTA2(S)-M2 (*92*) inducible promoter for 24 h with doxycycline [200 ng/mL] (Sigma, D9891). No respiratory repression of MCF10A or MB-MDA-231 cells was observed with 200 ng/mL, 1 μg/mL, or 2 μg/mL doxycycline. Previous studies have demonstrated that 24 h of 30 μg/mL doxycycline treatment can suppress mitochondrial respiration (*93*).

ATGAATTTACAACCAATTTTCTGGATTGGACTGATCAGTTCAGTTTGCTGTGTGTTTGCTCAAACAGATGGCGAGCAGAAGCTGATCAGCGAGGAGGACCTGGGCGAGCAGAAGCTGATCAGCGAGGAGGACCTGGGCGAGCAGAAGCTGATCAGCGAGGAGGACCTGGGCGGCGCCCAAACAGATGAAAATCGATGTTTAAAAGCAAATGCCAAATCATGTGGAGAATGTATACAAGCAGGGCCAAATTGTGGGTGGTGCACAAATTCAACATTTTTACAGGAAGGAATGCCTACTTCTGCACGATGTGATGATTTAGAAGCCTTAAAAAAGAAGGGTTGCCCTCCAGATGACATAGAAAATCCCAGAGGCTCCAAAGATATAAAGAAAAATAAAAATGTAACCAACCGTAGCAAAGGAACAGCAGAGAAGCTCAAGCCAGAGGATATTACTCAGATCCAACCACAGCAGTTGGTTTTGCGATTAAGATCAGGGGAGCCACAGACATTTACATTAAAATTCAAGAGAGCTGAAGACTATCCCATTGACCTCTACTACCTTATGGACCTGTCTTACTCAATGAAAGACGATTTGGAGAATGTAAAAAGTCTTGGAACAGATCTGATGAATGAAATGAGGAGGATTACTTCGGACTTCAGAATTGGATTTGGCTCATTTGTGGAAAAGACTGTGATGCCTTACATTAGCACAACACCAGCTAAGCTCAGGAACCCTTGCACAAGTGAACAGAACTGCACCAGCCCATTTAGCTACAAAAATGTGCTCAGTCTTACTAATAAAGGAGAAGTATTTAATGAACTTGTTGGAAAACAGCGCATATCTGGAAATTTGGATTCTCCAGAAGGTGGTTTCGATGCCATCATGCAAGTTGCAGTTTGTGGATCACTGATTGGCTGGAGGAATGTTACACGGCTGCTGGTGTTTTCCACAGATGCCGGGTTTCACTTTGCTGGAGATGGGAAACTTGGTGGCATTGTTTTACCAAATGATGGACAATGTCACCTGGAAAATAATATGTACACAATGAGCCATTATTATGATTATCCTTCTATTGCTCACCTTGTCCAGAAACTGAGTGAAAATAATATTCAGACAATTTTTGCAGTTACTGAAGAATTTCAGCCTGTTTACAAGGAGCTGAAAAACTTGATCCCTAAGTCAGCAGTAGGAACATTATCTGCAAATTCTAGCAATGTAATTCAGTTGATCATTGATGCATACAATTCCCTTTCCTCAGAAGTCATTTTGGAAAACGGCAAATTGTCAGAAGGAGTAACAATAAGTTACAAATCTTACTGCAAGAACGGGGTGAATGGAACAGGGGAAAATGGAAGAAAATGTTCCAATATTTCCATTGGAGATGAGGTTCAATTTGAAATTAGCATAACTTCAAATAAGTGTCCAAAAAAGGATTCTGACAGCTTTAAAATTAGGCCTCTGGGCTTTACGGAGGAAGTAGAGGTTATTCTTCAGTACATCTGTGAATGTGAATGCCAAAGCGAAGGCATCCCTGAAAGTCCCAAGTGTCATGAAGGAAATGGGACATTTGAGTGTGGCGCGTGCAGGTGCAATGAAGGGCGTGTTGGTAGACATTGTGAATGCAGCACAGATGAAGTTAACAGTGAAGACATGGATGCTTACTGCAGGAAAGAAAACAGTTCAGAAATCTGCAGTAACAATGGAGAGTGCGTCTGCGGACAGTGTGTTTGTAGGAAGAGGGATAATACAAATGAAATTTATTCTGGCAAATTCTGCGAGTGTGATAATTTCAACTGTGATAGATCCAATGGCTTAATTTGTGGAGGAAATGGTGTTTGCAAGTGTCGTGTGTGTGAGTGCAACCCCAACTACACTGGCAGTGCATGTGACTGTTCTTTGGATACTAGTACTTGTGAAGCCAGCAACGGACAGATCTGCAATGGCCGGGGCATCTGCGAGTGTGGTGTCTGTAAGTGTACAGATCCGAAGTTTCAAGGGCAAACGTGTGAGATGTGTCAGACCTGCCTTGGTGTCTGTGCTGAGCATAAAGAATGTGTTCAGTGCAGAGCCTTCAATAAAGGAGAAAAGAAAGACACATGCACACAGGAATGTTCCTATTTTAACATTACCAAGGTAGAAAGTCGGGACAAATTACCCCAGCCGGTCCAACCTGATCCTGTGTCCCA TTGTAAGGAGAAGGATGTTGACGACTGTTGGTTCTATTTTACGTATTCAGTGAATGGGAACAACGAGGTCATGGTTCATGTTGTGGAGAATCCAGAGTGTCCCACTGGTCCA GACATCATTCCAATTGTAGCTGGTGTT**<u>AAC</u>**GCTGGAATTGTTCTTATTGGCCTTGCA TTACTGCTGATATGGAAGCTTTTAATGATAATTCATGACAGAAGGGAGTTTGCTAAATTTGAAAAGGAGAAAATGAATGCCAAATGGGACACGGGTGAAAATCCTATTTATAAGAGTGCCGTAACAACTGTGGTCAATCCGAAGTATGAGGGAAAATGA

### WT β1 integrin

Weaver lab generated (Dr. Jonathan Lakins) puromycin lentiviral transfer vector expressing 3x myc tagged β1 integrin using the tetracycline rtTA2(S)-M2 (*92*) inducible promoter for 24 h with doxycycline [200 ng/mL] (Sigma, D9891).

ATGAATTTACAACCAATTTTCTGGATTGGACTGATCAGTTCAGTTTGCTGTGTGTTTGCTCAAACAGATGGCGAGCAGAAGCTGATCAGCGAGGAGGACCTGGGCGAGCAGAAGCTGATCAGCGAGGAGGACCTGGGCGAGCAGAAGCTGATCAGCGAGGAGGACCTGGGCGGCGCCCAAACAGATGAAAATCGATGTTTAAAAGCAAATGCCAAATCATGTGGAGAATGTATACAAGCAGGGCCAAATTGTGGGTGGTGCACAAATTCAACATTTTTACAGGAAGGAATGCCTACTTCTGCACGATGTGATGATTTAGAAGCCTTAAAAAAGAAGGGTTGCCCTCCAGATGACATAGAAAATCCCAGAGGCTCCAAAGATATAAAGAAAAATAAAAATGTAACCAACCGTAGCAAAGGAACAGCAGAGAAGCTCAAGCCAGAGGATATTACTCAGATCCAACCACAGCAGTTGGTTTTGCGATTAAGATCAGGGGAGCCACAGACATTTACATTAAAATTCAAGAGAGCTGAAGACTATCCCATTGACCTCTACTACCTTATGGACCTGTCTTACTCAATGAAAGACGATTTGGAGAATGTAAAAAGTCTTGGAACAGATCTGATGAATGAAATGAGGAGGATTACTTCGGACTTCAGAATTGGATTTGGCTCATTTGTGGAAAAGACTGTGATGCCTTACATTAGCACAACACCAGCTAAGCTCAGGAACCCTTGCACAAGTGAACAGAACTGCACCAGCCCATTTAGCTACAAAAATGTGCTCAGTCTTACTAATAAAGGAGAAGTATTTAATGAACTTGTTGGAAAACAGCGCATATCTGGAAATTTGGATTCTCCAGAAGGTGGTTTCGATGCCATCATGCAAGTTGCAGTTTGTGGATCACTGATTGGCTGGAGGAATGTTACACGGCTGCTGGTGTTTTCCACAGATGCCGGGTTTCACTTTGCTGGAGATGGGAAACTTGGTGGCATTGTTTTACCAAATGATGGACAATGTCACCTGGAAAATAATATGTACACAATGAGCCATTATTATGATTATCCTTCTATTGCTCACCTTGTCCAGAAACTGAGTGAAAATAATATTCAGACAATTTTTGCAGTTACTGAAGAATTTCAGCCTGTTTACAAGGAGCTGAAAAACTTGATCCCTAAGTCAGCAGTAGGAACATTATCTGCAAATTCTAGCAATGTAATTCAGTTGATCATTGATGCATACAATTCCCTTTCCTCAGAAGTCATTTTGGAAAACGGCAAATTGTCAGAAGGAGTAACAATAAGTTACAAATCTTACTGCAAGAACGGGGTGAATGGAACAGGGGAAAATGGAAGAAAATGTTCCAATATTTCCATTGGAGATGAGGTTCAATTTGAAATTAGCATAACTTCAAATAAGTGTCCAAAAAAGGATTCTGACAGCTTTAAAATTAGGCCTCTGGGCTTTACGGAGGAAGTAGAGGTTATTCTTCAGTACATCTGTGAATGTGAATGCCAAAGCGAAGGCATCCCTGAAAGTCCCAAGTGTCATGAAGGAAATGGGACATTTGAGTGTGGCGCGTGCAGGTGCAATGAAGGGCGTGTTGGTAGACATTGTGAATGCAGCACAGATGAAGTTAACAGTGAAGACATGGATGCTTACTGCAGGAAAGAAAACAGTTCAGAAATCTGCAGTAACAATGGAGAGTGCGTCTGCGGACAGTGTGTTTGTAGGAAGAGGGATAATACAAATGAAATTTATTCTGGCAAATTCTGCGAGTGTGATAATTTCAACT GTGATAGATCCAATGGCTTAATTTGTGGAGGAAATGGTGTTTGCAAGTGTCGTGTGTGTGAGTGCAACCCCAACTACACTGGCAGTGCATGTGACTGTTCTTTGGATACTAGTACTTGTGAAGCCAGCAACGGACAGATCTGCAATGGCCGGGGCATCTGCGAGTGTGGTGTCTGTAAGTGTACAGATCCGAAGTTTCAAGGGCAAACGTGTGAGATGTGTCAGACCTGCCTTGGTGTCTGTGCTGAGCATAAAGAATGTGTTCAGTGCAGAGCCTTCAATAAAGGAGAAAAGAAAGACACATGCACACAGGAATGTTCCTATTTTAACATTACCAAGGTAGAAAGTCGGGACAAATTACCCCAGCCGGTCCAACCTGATCCTGTGTCCCATTGTAAGGAGAAGGATGTTGACGACTGTTGGTTCTATTTTACGTATTCAGTGAATGGGAACAACGAGGTCATGGTTCATGTTGTGGAGAATCCAGAGTGTCCCACTGGTCCAGACATCATTCCAATTGTAGCTGGTGTGGTTGCTGGAATTGTTCTTATTGGCCTTGCATTACTGCTGATATGGAAGCTTTTAATGATAATTCATGACAGAAGGGAGTTTGCTAAATTTGAAAAGGAGAAAATGAATGCCAAATGGGACACGGGTGAAAATCCTATTTATAAGAGTGCCGTAACAACTGTGGTCAATCCGAAGTATGAGGGAAAATGA

### In vitro respirometry

Mitochondrial stress tests were performed with a Seahorse XF24e cellular respirometer on non-permeablized cells at ~ 96% confluence (100k cells/well) in V7 microplates, with XF assay medium supplemented with 1 mM pyruvate (Gibco), 2 mM glutamine (Gibco),and 5 or 25 mM glucose (Sigma) at pH 7.4 and sequential additions via injection ports of Oligomycin [1 μM final], FCCP [1 μM final], and Antimycin A/Rotenone [1 μM final] during respirometry (concentrated stock solutions solubilized in 100% ethanol [2.5 mM] for mitochondrial stress test compounds). OCR values presented with non-mitochondrial oxygen consumption deducted.

### Atomic force microscopy

AFM and analyses were performed using an MFP3D-BIO inverted optical atomic force microscope mounted on a Nikon TE2000-U inverted fluorescence microscope (Asylum Research). 100k cells were seeded onto fibronectin coated 15 mm^2^ coverslips and cultured for 24 h. Coverslips were anchored with permanent adhesive dots (Scotch, 00051141908113) to a glass slide that was then magnet-anchored to the stage of the microscope. All samples were measured in media with contact mode using Novascan cantilevers (5 μm radius, Probe 58, *k* = 0.06 N per m), which were calibrated using the thermal tune method. 36 force measurements were collected over a 250 × 250 μm grid per sample. The resulting force data were converted to elastic modulus values using the Hertz Model program (tissue samples were assumed to be noncompressible, and a Poisson’s ratio of 0.5 was used in the calculation of theYoung’s elastic modulus values) in IgorPro v.6.22, supplied by Asylum Research

### Knockdown of HSF1

pLKO.1 puro (Addgene #8453) was modified to carry:

Scr insert: 5’-CAACAAGATGAAGAGCACCAACTCGAGTTGGTGCTCTTCATCTTGTTGTTTTT, shRNA HSF1-1 TRCN0000007481 (HSF1):
5’-CCGGGCAGGTTGTTCATAGTCAGAACTCGAGTTCTGACTATGAACAACCTGCTTTTT, shRNA HSF1-2 TRCN0000318652 (HSF1):
5’-CCGGGCACATTCCATGCCCAAGTATCTCGAGATACTTGGGCATGGAATGTGCTTTT T

### Constitutively active HSF1

CD510B-1_pCDH-CMV-MCS-EF1-Puro (SystemBio) vector was modified to carry the hHSF1ΔRD (Δ221–315) transgene (*94*) under the CMV promoter.

ATGGATCTGCCCGTGGGCCCCGGCGCGGCGGGGCCCAGCAACGTCCCGGCCTTCCTGACCAAGCTGTGGACCCTCGTGAGCGACCCGGACACCGACGCGCTCATCTGCTGGAGCCCGAGCGGGAACAGCTTCCACGTGTTCGACCAGGGCCAGTTTGCCAAGGAGGTGCTGCCCAAGTACTTCAAGCACAACAACATGGCCAGCTTCGTGCGGCAGCTCAACATGTATGGCTTCCGGAAAGTGGTCCACATCGAGCAGGGCGGCCTGGTCAAGCCAGAGAGAGACGACACGGAGTTCCAGCACCCATGCTTCCTGCGTGGCCAGGAGCAGCTCCTTGAGAACATCAAGAGGAAAGTGACCAGTGTGTCCACCCTGAAGAGTGAAGACATAAAGATCCGCCAGGACAGCGTCACCAAGCTGCTGACGGACGTGCAGCTGATGAAGGGGAAGCAGGAGTGCATGGACTCCAAGCTCCTGGCCATGAAGCATGAGAATGAGGCTCTGTGGCGGGAGGTGGCCAGCCTTCGGCAGAAGCATGCCCAGCAACAGAAAGTCGTCAACAAGCTCATTCAGTTCCTGATCTCACTGGTGCAGTCAAACCGGATCCTGGGGGTGAAGAGAAAGATCCCCCTGATGCTGAACGACAGTGGCTCAGCACATGGGCGCCCATCTTCCGTGGACACCCTCTTGTCCCCGACCGCCCTCATTGACTCCATCCTGCGGGAGAGTGAACCTGCCCCCGCCTCCGTCACAGCCCTCACGGACGCCAGGGGCCACACGGACACCGAGGGCCGGCCTCCCTCCCCCCCGCCCACCTCCACCCCTGAAAAGTGCCTCAGCGTAGCCTGCCTGGACAAGAATGAGCTCAGTGACCACTTGGATGCTATGGACTCCAACGAGGATAACCTGCAGACCATGCTGAGCAGCCACGGCTTCAGCGTGGACACCAGTGCCCTGCTGGACCTGTTCAGCCCCTCGGTGACCGTGCCCGACATGAGCCTGCCTGACCTTGACAGCAGCCTGGCCAGTATCCAAGAGCTCCTGTCTCCCCAGGAGCCCCCCAGGCCTCCCGAGGCAGAGAACAGCAGCCCGGATTCAGGGAAGCAGCTGGTGCACTACACAGCGCAGCCGCTGTTCCTGCTGGACCCCGGCTCCGTGGACACCGGGAGCAACGACCTGCCGGTGCTGTTTGAGCTGGGAGAGGGCTCCTACTTCTCCGAAGGGGACGGCTTCGCCGAGGACCCCACCATCTCCCTGCTGACAGGCTCGGAGCCTCCCAAAGCCAAGGACCCCACTGTCTCCTAG

### SLC9A1 KO

MCF10A cells were transfected v*ia* PEI (https://www.addgene.org/protocols/transfection/) with pSpCas9(BB)-2A-GFP (PX458) - (Addgene #48138) carrying sgRNA for hSLC9A1 KO 5’-GTTTGCCAACTACGAACACG (SLC9A1:HGLibA_45399) and H^+^-suicide selected (*95*) four separate times to isolate SCL9A1 KOs. ~15% of the cells survived the first H^+^-suicide selection, ~90% survived the subsequent 4 sections.

### CRISPR-I (YME1L1)

MCF10A cells were transfected with a lentiviral vector expressing dCas9-KRAB under the EF1a promoter with blasticidin resistance (EF1a-dCas9-KRAB-Blast), a gift from Dr. Michal McManus. A second lentiviral guide vector with puromycin resistance was used to express the gRNAs for YME1L1: (1) GCAGTAGCTGTAGGAAGGGG, (2) CTCCTCCCAGAAACGGAAAA designed with the Broad institute GPP sgRNA Designer (https://portals.broadinstitute.org/gpp/public/analysis-tools/sgrna-design-crisprai?mechanism=CRISPRi). All sequence inserts were verified and KD was assessed via western blotting.

### C. elegans strains and maintenance

All *C. elegans* strains are derivatives of the Bristol N2 strain from Caenorhabditis Genetics Center (CGC) and are listed below. All worms are maintained at 15 °C on standard nematode growth media (NGM) plates and fed OP50 *E. coli* B bacteria and are maintained for a maximum of 20-25 generations (weeks). For all experiments, worms are synchronized using a standard bleaching protocol by degrading carcasses with bleach solution (1.8% sodium hypochlorite, 0.375M KOH), then washing eggs four times with M9 solution (22 mM KH_2_PO_4_ monobasic, 42.3 mM Na_2_HPO_4_, 85.6 mM NaCl, 1 mM MgSO_4_), followed by L1 arresting synchronization, achieved by floating eggs in M9 overnight in a 20 °C incubator on a rotator for a maximum of 16 hours. L1s are plated on RNAi bacteria (NGM + 1 μM IPTG and 100 μg/mL carbenicillin; HT115 *E. coli* K strain containing pL4440 vector control or pL4440 with RNAi of interest) until the desired stages of adulthood. All RNAi constructs were isolated from the Vidal library and verified sequences are available below.

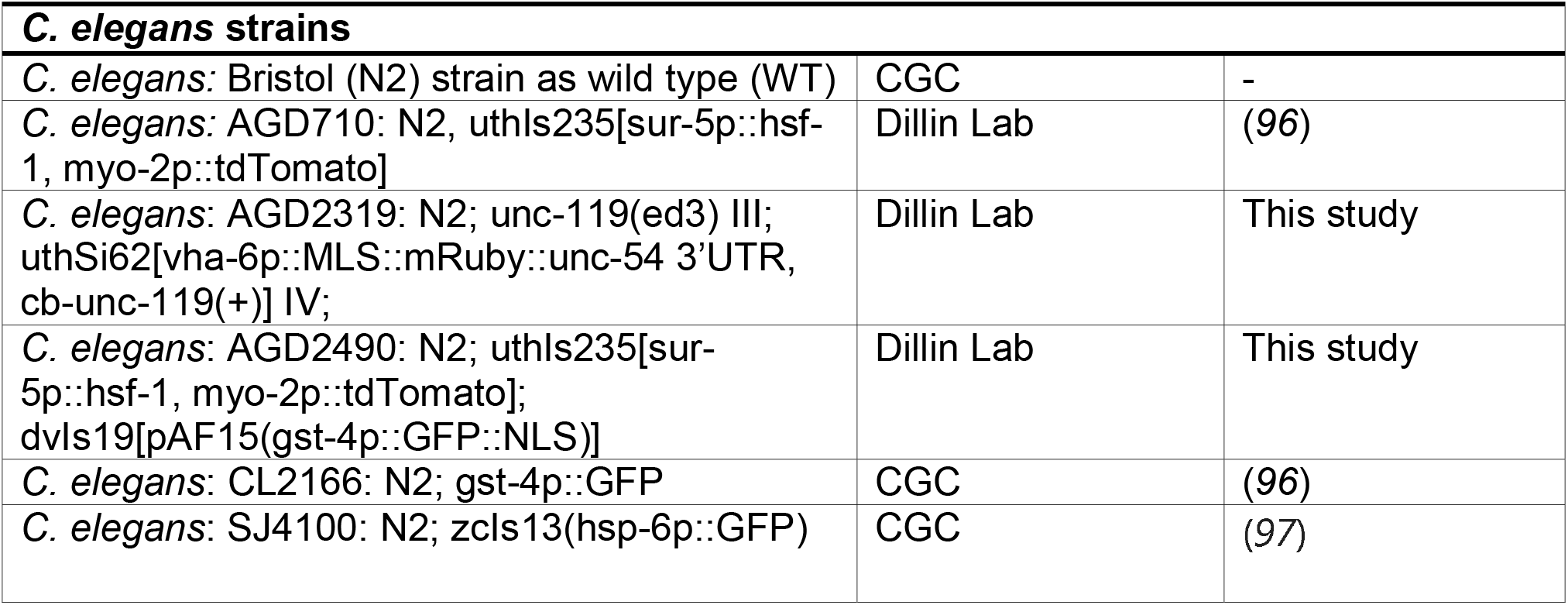

#### hsf-1

*CGTCA GCGCGCCCGTA CCGGCA CA TCAAA TCCA TTTTCCGGGTACTGTTGCTCA TT ATCAAACAACAAATCCTCGGCTCCA TCATAA TTCGACGTCTCCGGAGCA TTTTCAA GAGCAA GCTGTCTGAGTGGA TCCTCGGATCCTTCTTCA TCATCA TCCAACGGTACA TTATTCCCAAAATCATCCCAATTATGATTACTGACTAAATCTCTGAAACTCTCCAATGAAGTATCAGTTCCAGTGAAATATTCTTGAAGTTCTTGAGATAATTGACGATCAAATGATGGAGAGAGTCCGAGAGTTGGAGAATATAGATTTTGATGAGGATCTGCGTTGGTGGATGAGGTGGAAGTCGTTGGATGATGCTGATCTTCTATTGCCATTAGCTTCTGATGCGGTTGAAGGTATTGATGAGATGGTTGATATGGAATCATTGAAGGATCTGAAGGCATGAAGCCACTGTAATTGTTCACAAATCCTCCCGAATAGTCTTGTTGCGGCTGAAAATTTCGAATTTTTAGAC*

#### nhx-2

*GTCAACGAAGTTCTTTTTATTGCCGTGTTCGGAGAATCTCTATTGAACGACGGAGTTGCTGTTGTACTTTATCGAATGTTCTTGACCTTCTCTGAAATTGGAACTGAGAATCTGATAACATCTGATTATATCAATGGAGGTGTTTCTTTCCTGGTTGTTGCATTTGGTGGAATTGGAATTGGTCTTCTTTTTGCATTTTTGACAAGTCTTGTTACAAGATTTGCTAGAGATGAAGAGGTCAAAGTGCTCAATTCTGTATTTATTCTCATTCTTCCGTACACTTGTTATCTTTGTGGAGAACTTTTCGGTCTTTCAAGTATTATGGC*

#### ina-1

*AACATCAAATGGATACGTTTCAAATGTCGGCGAAAAGGATTATCTGGACTTGACATTCACTGTGGAAAACAAGAAAGAAAAGGCTTATCAAGCGAATTTTTATCTAGAATATAATGAAGAAGAGCTTGAACTTCCACAAGTTCAAGGTTCCAAGAGAATGATTGCTGAAACAATTGGAAAGAATATTGTGCATTTGCCACTTGGAAATCCAATGAATGGAGCATCAAAACATCAATTTACGATCCAATTCAAATTGACTCGTGGAAGAACTGAAGGAATTGGAAAGGCACTCAAATTCATGGCACATGTCAATTCCACGTCACAAGAAACCGAGGAAGAGTTGAAAGATAATAAATGGGAAGCTGAAGTTCAGATTATCAAGAAGGCAGAGCTGGAGATCTATGGAATCAGTGACCCTGATAGAGTATTCTTTGGAGGAAAAGCAAGAGCAGAGTCTGAATTGGAATTGGAAGAAGATATTGGAACAATGGTTAGACATAACTATACAATTATTAATCATGGTCCATGGACTGTTCGAAATGTGGAAGCACACATTTCTTGGCCTTATCAACTCCGTTCTAGGTTTGGAAGAGGAAAGAATGCTCTCTATCTATTGGATGTACCGACTATTACAACAGAATTCACAGATGGAACAAGTGAAGTTAGAAAGTGTTTCATCAAACAACAGTACGAATATGTGAATCCTGCAGAAATTAAATTGAACACTAAATATAGTACTCAAGAGACTGCTCCACATCGGGTAGAGCATAGAATGAAAAGAGAAATTGATGAGGATGAGGAAGAACAATCAGATGATCTAGGAGCCGTGGAAGAGAATATTCCTTGGTTCTCAACAGCTAATTTCTGGAATCTTTTTGCAATTAAAGGAGGTGATGGACG*

#### pat-3

*GATATGGTTTGTGGAGTTTGTCGATGCAAAGGAGGAAATGTTGGAAAATATTGTGAATGTAATAGACCTGGAATGAGTACTGCTGCGCTCAATGAAAAATGCAAAAGAACTAACGAATCAGCAATCTGTGAGGGTCGTGGTGTGTGTAACTGTGGACGTTGTGAATGTAATCCACGTGCCAATCCAGAAGAACAAATCTCTGGAGAATTCTGTGAATGCGACAACTTCAATTGCCCACGACACGATCGTAAAATCTGTGCAGAACACGGTGAATGCAACTGTGGAAAGTGTATTTGTGCACCTGGATGGACTGGAAGAGCCTGTGAATGCCCAATTTCAACTGATTCATGCCTCTCTGCAAATGGAAAAATCTGTAATGGAAAGGGTGAATGTATTTGTGGAAGATGTCGATGCTTCGATTCGCCCGACGGAAATCGATATTCGGGA GCGAAATGCGAAATTTGTCCGACGTGTCCGACGAAATGTGTGGAATACAAGAATTGTGTAATGTGCCAGCAATGGCAGACAGGGCCACTTAATGAGACCGCCTGTGATCAGTGTGAATTCAAAGTTATTCCTGTTGAGGAATTACCCAATCTCAACGAAACTACACCCTGCCAATTTGTGGATCCAGCTGATGATTGTACATTCTATTATCTCTACTATTACGATGAGGCCACAGATAATGCAACAGTCTGGGTCAGAAAACATAAAGATTGTCCTCCACCTGTCCCTGTGCTCGCAATTGTGCTCGGAGTCATTGCGGGTATCGTAATCCTCGGAATTCTTCTCTTGTTGCT*

#### skn-1

*TGTACACGGACAGCAATAATAGGAGCTTTGATGAAGTCAACCATCAGCATCAACAAGAACAAGATTTCAATGGCCAATCCAAATATGATTATCCACAATTCAACCGTCCAATGGGTCTCCGTTGGCGTGATGATCAACGGATGATGGAGTATTTCATGTCGAATGGTCCAGTAGAAACTGTTCCAGTTATGCCAATACTCACCGAGCATCCACCAGCATCTCCATTCGGTAGAGGACCATCTACAGAACGTCCAACCACATCATCTCGATACGAGTACAGTTCGCCTTCTCTCGAGGATATCGACTTGATTGATGTGCTATGGAGAAGTGATATTGCTGGAGAGAAGGGCACACGACAAGTGGCTCCTGCTGATCAGTACGAATGTGATTTGCAGACGTTGACAGAGAAATCGACAGTAGCG*

#### ymel-1

*TCGATTCAGTTGGCTCAAAACGTGTTTCGAATTCCATCCATCCATATGCAAATCAAACGATTAATCAACTTCTCAGGTAACAATTTGCTCAATTTTGTGCATTAAAAACTCATCTCCTGATGTTTTCAGTGAAATGGATGGCTTCACCCGTAACGAGGGAATCATTGTAATTGCCGCAACAAATCGTGTCGACGACCTC*

### C. elegans compound microscopy of mitochondria

Transgenic animals carrying vha-6p::MLS::mRuby (MLS was derived from *atp-1:* ATGTTGTCCAAACGCATTGTTACCGCTCTTAACACCGCCGTCAAGGTCCAAAATGCCGGAATCGCCACCACCGCCCGCGGA) were grown from L1 to desired stage of adulthood on standard RNAi plates as described above. Animals were aged by handpicking adults away from progeny using a pick daily until desired stage of adulthood. For imaging, adult worms are mounted on a glass slide in M9 solution, covered with a cover slip, and imaged immediately for a maximum of 10 minutes per slide. Animals were imaged on a Zeiss AxioObserver.Z1 microscope equipped with a lumencor sola light engine and a Zeiss axiocam 506 camera, driven by Zeiss ZenBlue software using a 63x/1.4 Plan Apochromat objective and a standard dsRed filter was used (Zeiss filter set 43).

### C. elegans Paraquat survival assay

Animals were grown to day 1 adulthood on standard RNAi plates as described above. 10 animals were picked into 75 μL of 100 mM paraquat solution prepared in M9 in a flatbottom 96-well plate. >8 wells are used per condition for a minimum of 80 animals per replicate. Animals were scored every 2 hours for death. Plates are tapped gently, and any trashing or bending movement is scored as alive. Paraquat survival assays are performed with the experimenter blinded to the strain conditions during scoring and are repeated a minimum of 3 replicates per experiment.

### C. elegans stereomicroscopy for fluorescent transcriptional reporters

Transgenic animals carrying *gst-4p::GFP* were grown on standard RNAi plates as described above until the L4 stage. L4 animals were washed off of plates using M9, centrifuged to pellet, and M9 was replaced with 50 mM paraquat prepared in M9. Animals were incubated rotating in a 20 °C incubator for two hours, and subsequently washed 2x with M9 solution. Animals were then plated on OP50 plates and recovered for 2 hours at 20 °C. For imaging, worms were picked onto a standard NGM plate containing 5 μL of 100 mM sodium azide to paralyze worms. Paralyzed worms were lined up, and imaged immediately on a Leica M250FA automated fluorescent stereomicroscope equipped with a Hamamatsu ORCA-ER camera, standard GFP filter, and driven by LAS-X software.

### C. elegans biosorter analysis

For large-scale quantification of fluorescent animals, a Union Biometrica complex object parameter analysis sorter (COPAS) was used (for full details, refer to (*98*). Briefly, to quantify signal of *gst-4p::GFP*, animals treated as described above were washed off plates using M9, and run through the COPAS biosort using a 488 nm light source. Integrated fluorescence intensity normalized to the time of flight is collected automatically on the COPAS software, and then normalized again to the extinction to correct for both worm length and worm thickness.

For JC-9 staining, day 1 adult animals were transferred to a plate containing JC-9-treated bacteria (OP50 bacteria were grown during mid-log phase for 4 hours in LB containing 50 μM JC-9 at 37 °C to incorporate JC-9 into bacteria; then bacteria were washed 2x with fresh LB to remove excess JC-9). Animals were grown on JC-9 bacteria for 2 hours at 20 °C to label. After labeling, worms were moved onto standard OP50 plates and grown for an additional 1 hour at 20 °C to remove excess JC-9 from the gut. Animals were then washed off plates and immediately run on a biosorter using a 488 and 561 nm light source. Worm profile data was collected, and run through an orientation and quantification algorithm, LAMPro (*98*). Briefly, integrated fluorescence intensity is measured throughout the entire profile of the worm and normalized to extinction throughout the length of the worm. Total integrated fluorescence of the entire worm was also calculated by normalizing to the time of flight and integrated extinction of the entire worm. JC-9 fluorescence at 515 nm was used to determine mitochondrial quantity, and the ratio of the fluorescence of 585 nm / 515 nm was used to determine mitochondrial membrane potential.

### Lifespan assay

Lifespan measurements were performed on solid NGM plates with RNAi bacteria. Worms were synchronized via bleaching/L1 arrested as described above. Adult animals were moved away from progeny by moving worms onto fresh RNAi plates every day until D7-10 when progeny were no longer visible. Animals were then scored every 1-2 days for death until all animals were scored. Animals with bagging vulval explosion, or other age-unrelated deaths were censored and removed from quantification.

### Lattice Light Sheet Microscopy

We used a Custom build lattice light sheet microscope (*28*) to image MCF10A culture on polyacrylamide gels. Polyacrylamide gels (PA-gels) were formed on 5 mm round cover glass (Warner Instruments), coated with fibronectin and seeded with ~ 1000 cells per gel. The samples were cultured for 24 h in MCF10A media prior to imaging in DMEM (5 mM glucose) without phenol red supplemented with 5% Fetal Bovine serum. Samples were illuminated by 561 nm diode laser (0.5W Coherent) or 639 nm diode laser (1W Coherent) using an excitation objective (Special Optics, 0.65 NA with a working distance of 3.74-mm) at 2% AOTF transmittance and output laser power of 100 mW. The measured powers at the back focal plane of the illumination objective were in the range of 0.15-0.2 mW. Order transfer functions were calculated by acquiring Point-spread functions using 200-nm TetraSpeck beads adhered freshly to 5-mm glass coverslips (Invitrogen T7280) for each excitation wavelength and each acquisition filter set. The LLSM was realigned before each experiment.

For illumination we displayed on the spatial light modulator (SLM) a Square lattice generated by an interference pattern of 59 bessels beams separated by 1.67 um and cropped to 0.22 with a 0.325 inner NA and 0.40 outer NA, or by a an interference pattern of 83 bessels beams separated by 1.23 um and cropped to 0.22 with a 0.44 inner NA and 0.55 outer NA The lattice light sheet was dithered 15-25 um to obtain an homogenous illumination with 5% of flyback time. Fluorescent signal was collected by a Nikon detection objective (CFI Apo LWD 25XW, 1.1 NA, 2-mm working distance (WD)), coupled with a 500 mm focal length tube lens (Thorlabs), a set of Semrock filters (BL02-561R-25, BLP01-647R-25, and NF03-405-488-561-635E-25), and a sCMOS camera (Hamamatsu Orca Flash 4.0 v2) with a 103 nm/pixel magnification.

Z-Stacks (Volumes) were acquired by moving the Z-piezo in scanning mode while leaving the lattice light sheet static. The slices of the stacks were taken with an interval of 100-235 nm (S-axis) through ranges of 30-35 um at 20-100 ms exposuretime with 0 - 60 seconds intervals between volumes.

Raw data was flash corrected (*99*) and deconvolved using an iterative Richardson-Lucy algorithm (*28*) on two graphics processing units (GPU) (NVIDIA, GeForce GTX TITAN 4 Gb RAM). Flash calibration, flash correction, channel registration, Order transfer function calculation and Image deconvolution were done using the LLSpy open software (*100*). Visualization of the images and volume inspection were done using Spimagine (Github/maweigert/spimagine) and Clear volume (*101*).

For the glucose shock experiment, MCF10A cells were first localized under the LLSM. During the first 30 seconds of the acquisition, the glucose concentration was raised to a final concentration of 25 mM. The glucose infusion caused misalignment of the lattice light sheet microscope which was corrected manually during the first acquisition volume.

### RNAseq

Total RNA was isolated using Trizol (Invitrogen), and RNAseq libraries (2 biological replicates per condition comprised of a pool of 4 PA-gel cultures each) prepared using KAPA mRNA HyperPrep Kit (Roche) and IDT dual indexed sequencing adaptors. Multiplexed libraries were sequenced on an Illumina HiSeq4000, and reads were aligned to the human genome (hg19) using RNA STAR (*102*). Aligned reads were counted using HOMER (*103*), and hierarchical clustering was performed using Cluster (*104*) and visualized with Java TreeView. Gene Ontology analysis was performed using Metascape (*105*).

### Mitochondrial ETC proteomics timsTOF

1 million cells were seeded on 50 mM^2^ varied stiffness ECM coated PA-gels cultured for 24 h. Cells were washed with sterile PBS once, then cells were detached with cold PBS and a cell scraper (rubber policeman). Cell pellets were then suspended in 100 μL urea lysis buffer (ULB: 8M urea, 100 mM Tris, and 75 mM NaCl at pH 8). Probe sonicated 5 times for 2 s on ice. Protein concentrations were determined *via* BCA, and 100 μg of each sample was alkylated and reduced for 1 h at ~22 °C (protected from light) *via* the addition of a 10X concentrated stock (ARB: 400 mM 2-Chloroacetamide and 100 mM Tris(2-carboxyethyl)phosphine (TCEP) dissolved in ULB) yielding a final concentration of 40 mM 2-Chloroacetamide and 10 mM TCEP. Samples were then diluted, with a Tris/NaCl buffer (100 mM Tris and 75 mM NaCl at pH 8) containing 1 μg LysC (Promega, Va11A) per sample, to yield a final concertation of 2M urea during a 4 h digestion at ~22 °C (protected from light). Following the LysC digestion, 2 μg of Trypsin (Pierce, 1862746) was added and allowed to react with the sample overnight at 37 °C. The samples were then acidified with Trifluoroacetic acid (TFA) [~1% final] to yield a sample pH of 2. Samples were then desalted on C18 tips (Nest Group), the eluant was lyophilized, and then resuspended in 4% formic acid, 3% acetonitrile at 200 fmol/μL concentration. For each MS analysis, 1 μL of sample was separated over a 25 cm column packed with 1.9 μm Reprosil C18 particles (Dr. Maisch HPLC GmbH) by a nanoElute HPLC (Bruker). Separation was performed at 50 °C at a flow rate of 400 μL/min by the following gradient in 0.1% formic acid: 2% to 17% acetonitrile from 0 to 60 min, followed by 17% to 28% acetonitrile from 60 to 105 min. The eluant was directed electrospray ionized into a Bruker timsTOF Pro mass spectrometer and data was collected using data-dependent PASEF acquisition (*106*). Database searching and extraction of MS1 peptide abundances was performed using the MaxQuant algorithm (*107*), and all peptide and protein identifications were filtered to a 1% false-discovery rate. Searches were performed against a protein database of the human proteome (downloaded from Uniprot on 3/21/2018). Lastly quality control analysis was performed via artMS (http://artms.org) and statistical testing was performed with MSstats (*108*).

### LC-MS/MS Deuterium Incorporation Proteomics

1 million cells were seeded on 50 mM^2^ varied stiffness PA-gels cultured for 24 h in 6% D_2_O culture media in a 5% CO_2_ incubator humidified with 5% D_2_O. D_2_O labeled cells were detached with cold PBS and a cell scraper (rubber policeman), pelleted with centrifugation, and mitochondrial fractions were isolated with the Mitochondria Isolation Kit for Cultured Cells (Thermo, 89874). Protein was isolated by flash freezing and sonication in PBS with 1 mM PMSF, 5 mM EDTA, and 1x Halt protease inhibitor (Thermo, 78440). Protein content was quantified via BCA (Pierce, 23225) and 100 μg of protein from each sample was trypsin (Pierce, 90057) digested overnight after reduction and alkylation with DTT, TFE, and iodoactamide (*109*). Trypsin-digested peptides were analyzed on a 6550 quadropole time of flight (Q-ToF) mass spectrometer equipped with Chip Cube nano ESI source (Agilent Technologies). High performance liquid chromatography (HPLC) separated the peptides using capillary and nano binary flow. Mobile phases were 95% acetonitrile/0.1% formic acid in LC-MS grade water. Peptides were eluted at 350 nL/minute flow rate with an 18 minute LC gradient. Each sample was analyzed once for protein/peptide identification in data-dependent MS/MS mode and once for peptide isotope analysis in MS mode. Acquired MS/MS spectra were extracted and searched using Spectrum Mill Proteomics Workbench software (Agilent Technologies) and a human protein database (www.uniprot.org). Search results were validated with a global false discovery rate of 1%. A filtered list of peptides was collapsed into a nonredundant peptide formula database containing peptide elemental composition, mass, and retention time. This was used to extract mass isotope abundances (M0-M3) of each peptide from MS-only acquisition files with Mass Hunter Qualitative Analysis software (Agilent Technologies). Mass isotopomer distribution analysis (MIDA) was used to calculate peptide elemental composition and curve-fit parameters for predicting peptide isotope enrichment based on precursor body water enrichment (p) and the number (n) of amino acid C-H positions per peptide actively incorporating hydrogen (H) and deuterium (D) from body water. Subsequent data handling was performed using python-based scripts, with input of precursor body water enrichment for each subject, to yield fractional synthesis rate (FSR) data at the protein level. FSR data were filtered to exclude protein measurements with fewer than 2 peptide isotope measurements per protein. Details of FSR calculations and data filtering criteria were described previously (Holmes, W.E., et al., 2015).

### LC-MS/MS Metabolomics

1 million cells were seeded on 50 mM^2^ varied stiffness ECM coated PA-gels cultured for 24 h. Cells were dissolved in 100% methanol doped with NEM [8 mM,1 mg/mL] (Sigma-Aldrich, cat. no. E1271) (*110*). Protein concentrations of the methanol extract was determined via BCA (Pierce, 23225) Data was normalized to 100 μg per sample and polar metabolites were extracted in a total volume of 275 μl of 40:40:20 (acetonitrile:methanol:water) with inclusion of internal standard d_3_N^15^-serine (Cambridge Isotope Laboratories, Inc. #DNLM-6863). Extracted samples were centrifuged at 10,000 × g for 10 min and an aliquot of the supernatant was injected onto LC/MS where metabolites were separated by liquid chromatography as previously described (*111*). Analysis was performed with an electrospray ionization (ESI) source on an Agilent 6430 QQQ LC-MS/MS (Agilent Technologies). The capillary voltage was set to 3.0 kV, and the fragmentor voltage was set to 100 V, the drying gas temperature was 350 °C, the drying gas flow rate was 10 L/min, and the nebulizer pressure was 35 PSI. Polar metabolites were identified by SRM of the transition from precursor to product ions at associated optimized collision energies and retention times as previously described (*112*). Quantification of metabolites was performed by integrating the area under the curve and then normalizing to internal standard values. All metabolite levels are expressed as relative abundances compared to the control group.

### Paraquat survival

10 mM paraquat (Acros Organics, 227320010) was dissolved into media and added to cell culture vessels and allowed to affect the cells for 24 h, experiments were always performed with a fresh suspension of paraquat. After 24 h of paraquat treatment cells were fixed with 4% PFA and stained for cleaved caspase 3 (Cell Signaling, 9661).

## Supplemental Videos

Video S1- Lattice Light Sheet Mitotracker Scan - Soft (400 Pa)

Video S2 - Lattice Light Sheet Mitotracker Scan - Stiff (60k Pa)

Video S3 - Lattice light Sheet Mitotracker Z-Stack Scan - Soft (400 Pa)

Video S4 - Lattice Light Sheet Mitotracker 3D Reconstruction - Soft (400 Pa)

Video S5 - Combined Stacks of Mitotracker - 6k Pa PA-gel - Initiates Toroid Formation

Video S6 - Video of Figure S1H

Video S7 - Figure S1H RED LUT

Video S8 - Lattice Light Sheet Mitotracker Scan - Stiff (60k Pa)

Video S9 - Lattice Light Sheet Mitotracker Scan - Stiff (60k Pa) with Mitotempo

**Figure S1:**
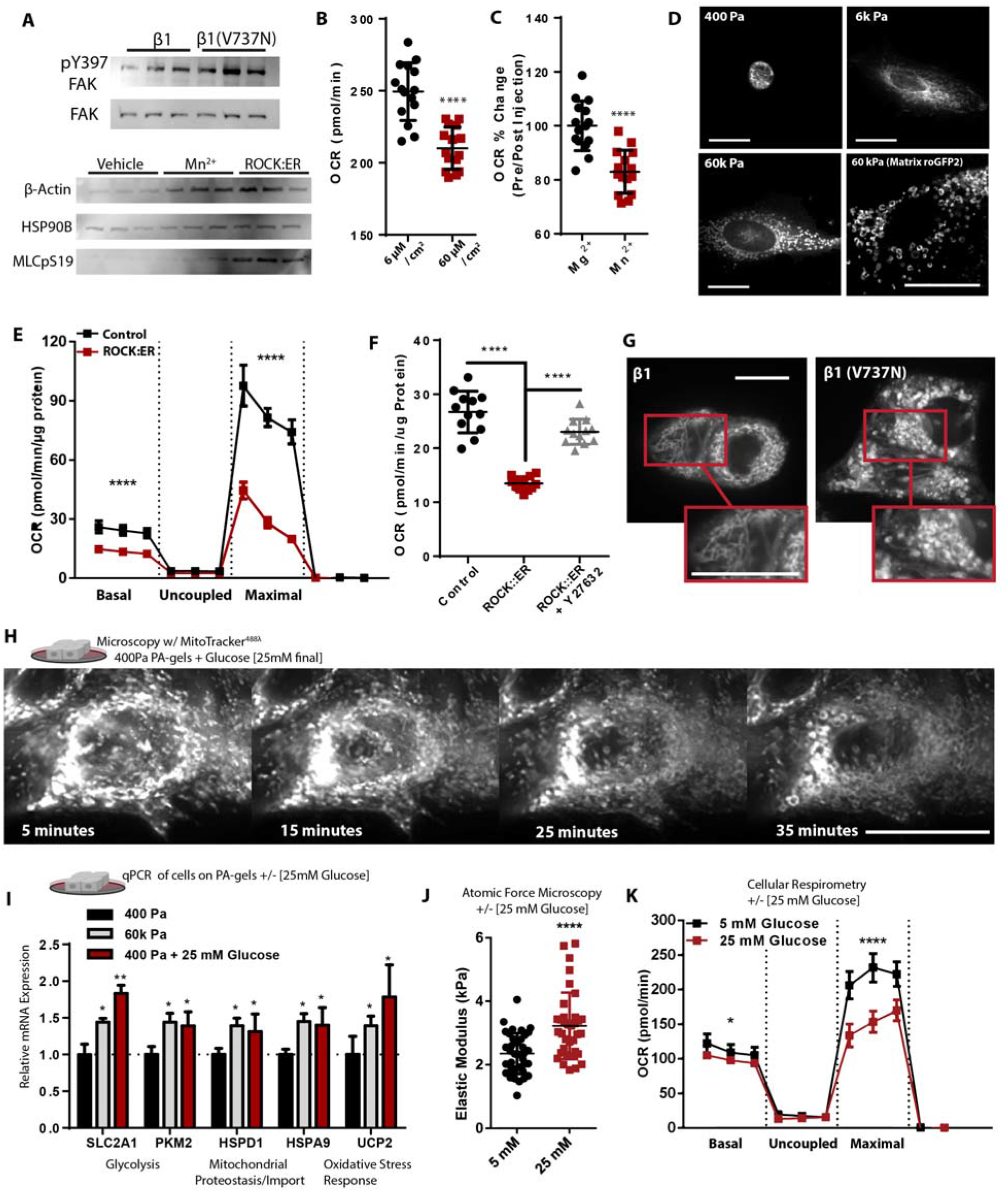
A. Western blot depicting relative protein abundance of focal adhesion kinase (FAK) and phosphorylated/active FAK-pY397 within 5 μg of total protein derived from lysates of β1-integrin or β1(V7373N) expressing cells (n=3 separate experiments). Western blot depicting relative protein abundance of β-actin, HSP90B, and ROCK phosphorylated myosin light chain (MLC-pS19) within 5 μg of total protein derived from lysates of ROCK:ER fusion protein expressing cells activated *via* 4HT [1 μM] treatment for 48 h or vehicle +/- 30 min of Mn^2^+ [1 μM]. (n=3 separate experiments). B. OCR of cells cultured for 24 h on varied fibronectin surface coating density (n=5 wells, 3 replicate measures). C. OCR 15 min before and a 15 minutes after the addition of Mn2+ or Mg2+ [1 μM] via injection port (n=5 wells, 3 replicate measures, repeated 3 times). D. Microscopy depicting mitochondrial network structure in PFA-fixed single cells cultured on soft-to-stiff fibronectin [6 μM/cm^2^] coated polyacrylamide hydrogels, mitotracker (deep red FM) [100 nM]. Mitochondrial matrix targeted roGFP2 expressing MEC cultured on stiff ECM for structure assessment of genetically encoded fluorophore relative to mitotracker staining (Scale Bar: 10 μm) E. Mitochondrial stress test of ROCK:ER fusion protein expressing cells activated *via* 4HT [1 μM] treatment for 48 h or vehicle (n=5 wells, 3 replicate measures, repeated 3 times). F. OCR of ROCK:ER fusion protein expressing cells activated *via* 4HT [1 μM] treatment for 48 h or vehicle, +/- y27632 [10 μM] (n=4 wells, 3 replicate measures, repeated 3 times). G. Confocal microscopy depicting mitochondrial network structure of β1-integrin or β1(V7373N) expressing PFA-fixed cells cultured on 400 Pa ECM for 24 h, stained with mitotracker (deep red FM) [100 nM]. (Scale Bar: 10 μm) H. Representative microscopy of the mitochondrial network structure in live cells cultured in 5 mM glucose on 400 Pa ECM with a lattice light sheet microscope, monitoring structural changes after infusion of glucose [200 mM], rendering a final concentration of 25 mM glucose, mitotracker (Green FM) [100 nM]. I. Relative gene expression of MECs grown of soft (400 PA) or stiff (60k Pa) ECM +/- glucose [5 or 25 mM] media normalized to low glucose media [5 mM], qPCR-ΔΔCT (housekeeping gene: 18s) (n=5 or 4 separate experiments). J. Cellular elasticity of MECs exposed to 24 h glucose [5 or 25 mM] on fibronectin [6 μM/cm^2^] coated glass coverslips, atomic force microscopy (n= 36 indentations across 3 discrete samples). K. OCR of cells in either 5 or 25mM Seahorse CF assay media, previously cultured in 25 mM glucose MCF10A media (n=5 wells, 3 replicate measures, repeated 3 times). *Data shown represent ± SEM. *P < 0.05, **P < 0.01, ***P < 0.005, < 0.0001 via two-tailed unpaired Student t test*.

**Figure S2:**
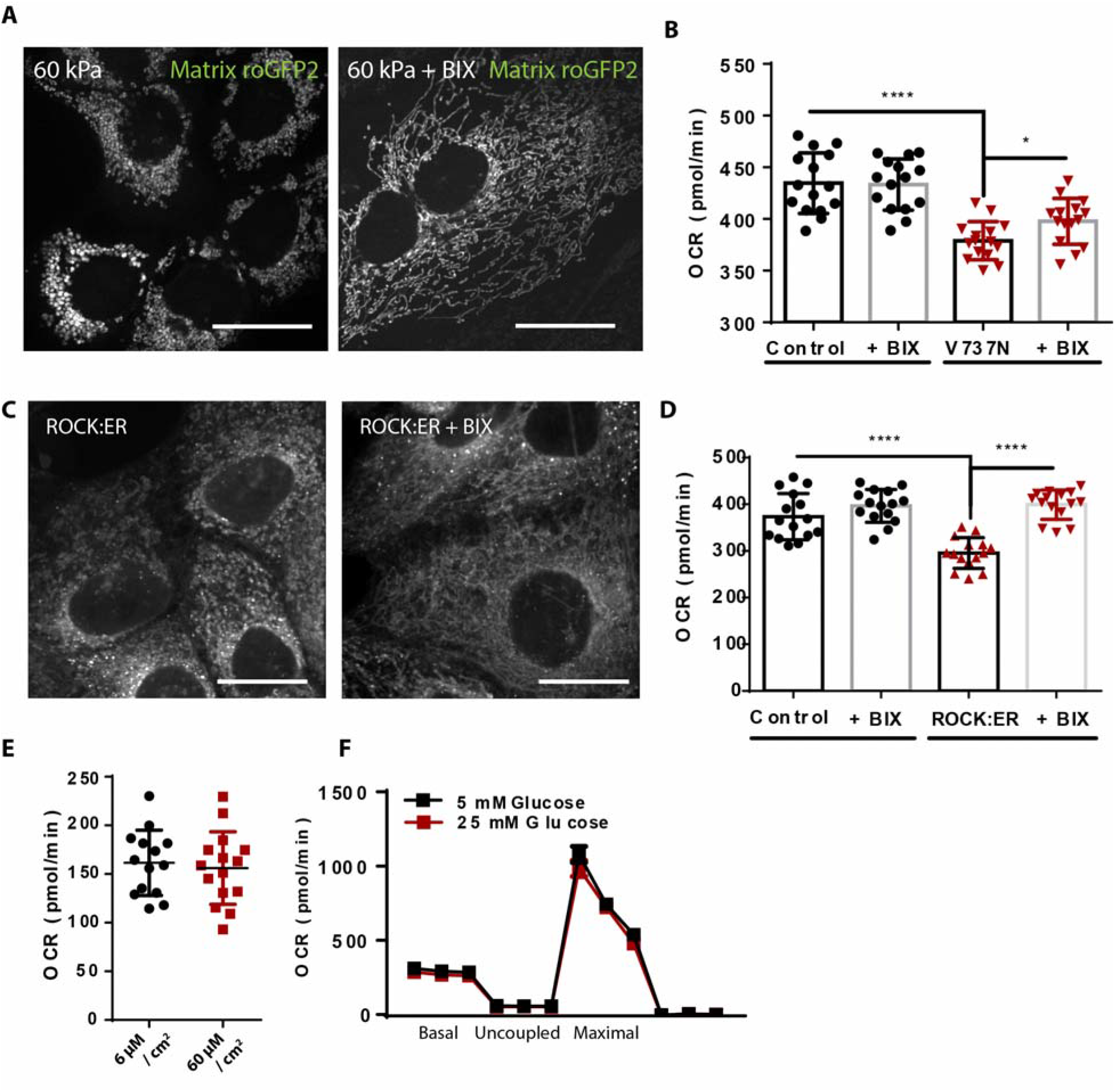
A. Confocal microscopy depicting mitochondrial network structure in cells expressing mitochondrial matrix targeted roGFP2 cultured on 60k Pa PA-gel surfaces +/- BIX [500 nM]. (Scale Bar: 10 μm) B. OCR of β1(V7373N) expressing cells via tetracycline inducible promoter for 24 h with doxycycline [200 ng/mL] +/- BIX [500 nM], (n=5 wells, 3 replicate measures, repeated 3 times). E. Confocal microscopy depicting mitochondrial network structure in ROCK:ER fusion protein expressing cells activated via 4HT [1 μM] treatment for 48 h or vehicle +/- BIX [500 nM]. (Scale Bar: 10 μm) D. OCR of ROCK:ER fusion protein expressing cells activated *via* 4HT [1 μM] treatment for 48 h or vehicle, +/- BIX [500 nM] (n=5 wells, 3 replicate measures, repeated 3 times). E. Oxygen consumption rate (OCR) of SLC9A1 KO cells in response to glucose [5 or 25 mM] (n=5 wells, 3 replicate measures). F. Mitochondrial stress test of SLC9A1 KO cells in response to 24 h of varied fibronectin surface coating density (n=5 wells, 3 replicate measures). *Data shown represent ± SEM. *P < 0.05, **P < 0.01, ***P < 0.005, < 0.0001 via twotailed unpaired Student t test (E) or one-way ANOVA with Tukey test for multiple comparisons (B and D*).

**Figure S3:**
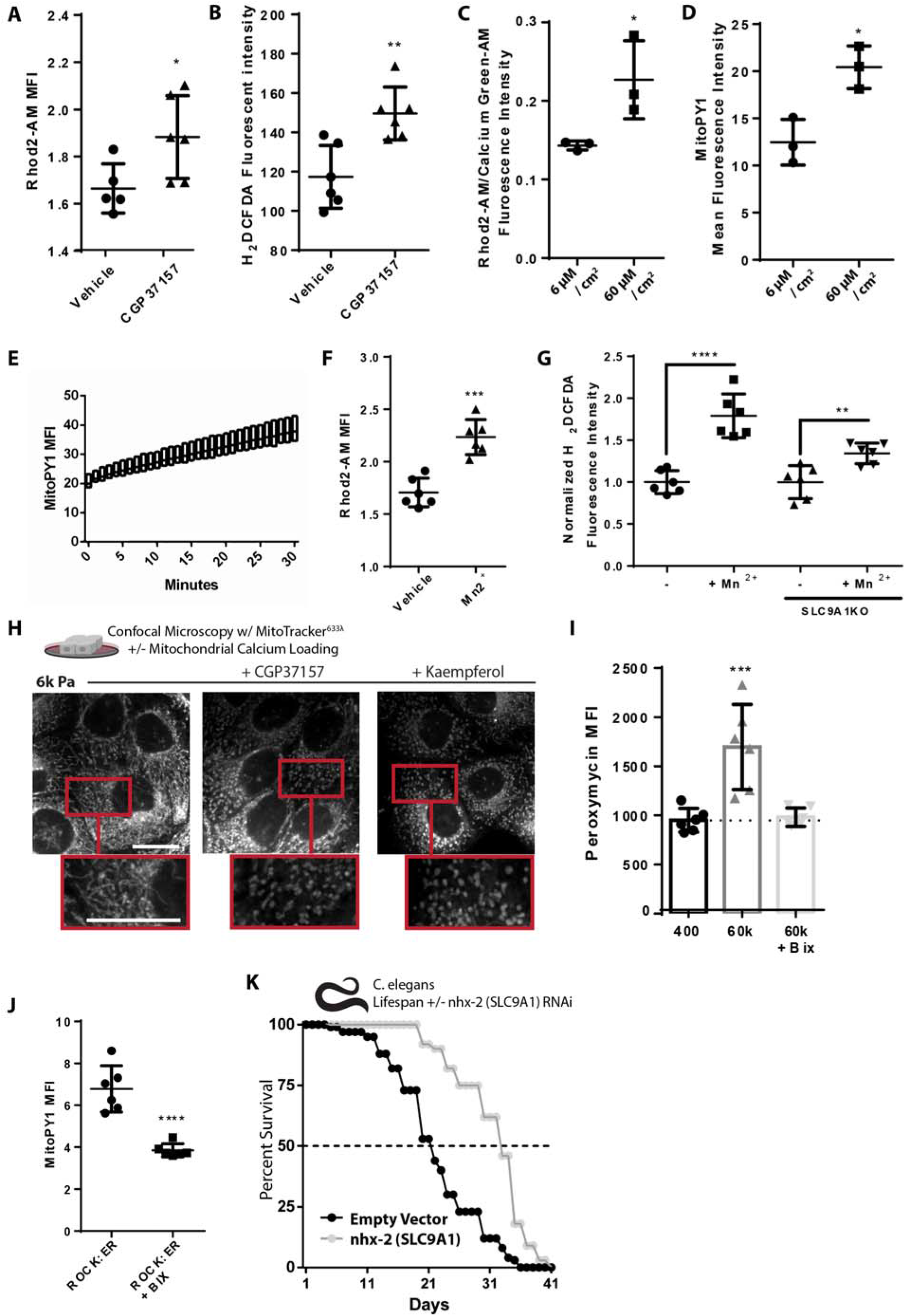
A. Mitochondrial calcium content of cells treated with CGP37157 [1 μM] for 1 h, Rhod2-AM [2 μM] (n=6 wells, repeated 2 times). B. Oxidative stress indicator intensity of cells treated with CGP37157 [1 μM] for 1 h, measured with 2’,7’-dichlorodihydrofluorescein diacetate (H2DCFDA) [2 μM] (n=6 wells, repeated 2 times). C. Calcium content of MCF10A cells cultured for 24 h of varied fibronectin surface coating density, treated with Rhod2-AM [2 μM] (mitochondrial) and Calcium Green-1-AM [2 μM] (intracellular). (n=4 replicates, repeated 2 times). D. Mitochondrial H_2_O_2_ production of MCF10A cells cultured for 24 h of varied fibronectin surface coating density, MitoPy1 [5 μM] (n=4 replicates, repeated 2 times). E. Mitochondrial H_2_O_2_ production of MCF10A cells in response to increased glucose concentration [25 mM], measured over 30 minutes after glucose concentration was increased from 5 to 25 mM, MitoPy1 [5 μM] (n=6, repeated 2 times). F. Mitochondrial calcium content of cells treated with Mn2+ [1 μM] for 30 min, Rhod2-AM [2 μM] (n=6 wells, repeated 2 times). G. GOxidative stress indicator intensity of WT or SLC9A1 KO cells treated with Mn2+ [1 μM] for 30 min, measured with 2’,7’-dichlorodihydrofluorescein diacetate (H2DCFDA) [2 μM] (n=6 wells, repeated 2 times). H. Confocal microscopy depicting mitochondrial network structure of PFA-fixed cells cultured on 6k Pa surfaces treated with CGP37157 [1 μM], kaempferol [10 μM], or vehicle for 1 h, mitotracker (deep red FM) [100 nM]. I. H_2_O_2_ production of cells cultured on 400 Pa or 60k Pa ECM +/- BIX [500 nM] for 24 h measured with peroxymycin fluorescent intensity (n=6 separate samples). J. Mitochondrial H2O2 production of ROCK:ER fusion protein expressing cells activated via 4HT [1 μM] treatment for 48 h +/- BIX [500nM] or vehicle for 24 h and then MitoPy1 [5 μM] (n=6 wells, repeated 2 times). K. Lifespan of *C. elegans* grown on nhx-2 (SLC9A1 orthologue) or empty vector RNAi, (n=120 animals, repeated 3 times) *Data shown represent ± SEM. *P < 0.05, **P < 0.01, ***P < 0.005, < 0.0001 via twotailed unpaired Student t test (A-D, F, G, and J) or one-way ANOVA with Tukey test for multiple comparisons (I*).

**Figure S4:**
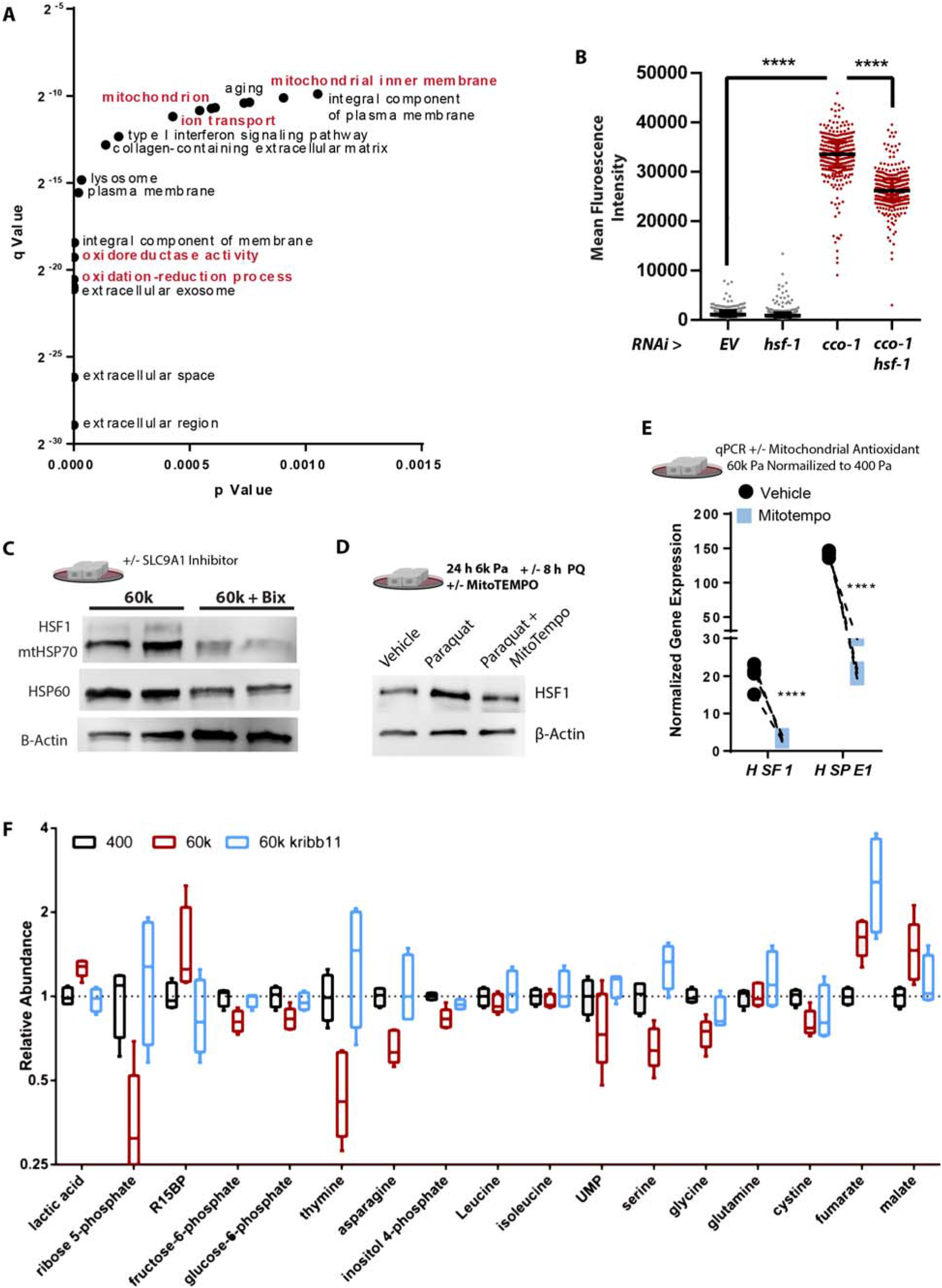
A. Gene ontology of categories more highly represented in cells cultured on 400 Pa ECM than 60k Pa ECM (n=2 duplicate libraries of 3 biological replicates, ~ 10 million reads per library). B. *hsp-6::gfp* reporter fluorescent intensity of *C.elegans* measured with a large particle cytometer, +/- *cco-1* RNAi +/- HSF1 RNAi (1:1 ratio) (n=213, 263, 295, and 285 animals in order left to right), repeated 3 times. C. Western blot depicting relative protein abundance of 60k Pa ECM +/- 24 h Bix [500 nM] treatment (n=2 two biological replicates shown, repeated 3 times). D. Western blot depicting relative protein abundance of HSF1 and β-actin within 5 μg of total protein derived from lysates of cells cultured 6k Pa ECM +/- MitoTempo treatment for 24 h +/- paraquat [10 mM] treatment for 4 additional hours. E. Relative HSF1 and HSPE1 (HSF1 target gene) mRNA expression of cells grown 60k Pa ECM normalized to 400 Pa ECM, qPCR-ΔΔCT (housekeeping gene: 18s) (n= 4 separate experiments). F. Selection of metabolites sensitive to stiffness or KRIBB11 treatment (n=5 biological replicates), related to Figure 4I). *Data shown represent ± SEM. *P < 0.05, **P < 0.01, ***P < 0.005, < 0.0001 via twotailed unpaired Student t test (E) or one-way ANOVA with Tukey test for multiple comparisons (B*).

**Figure S5:**
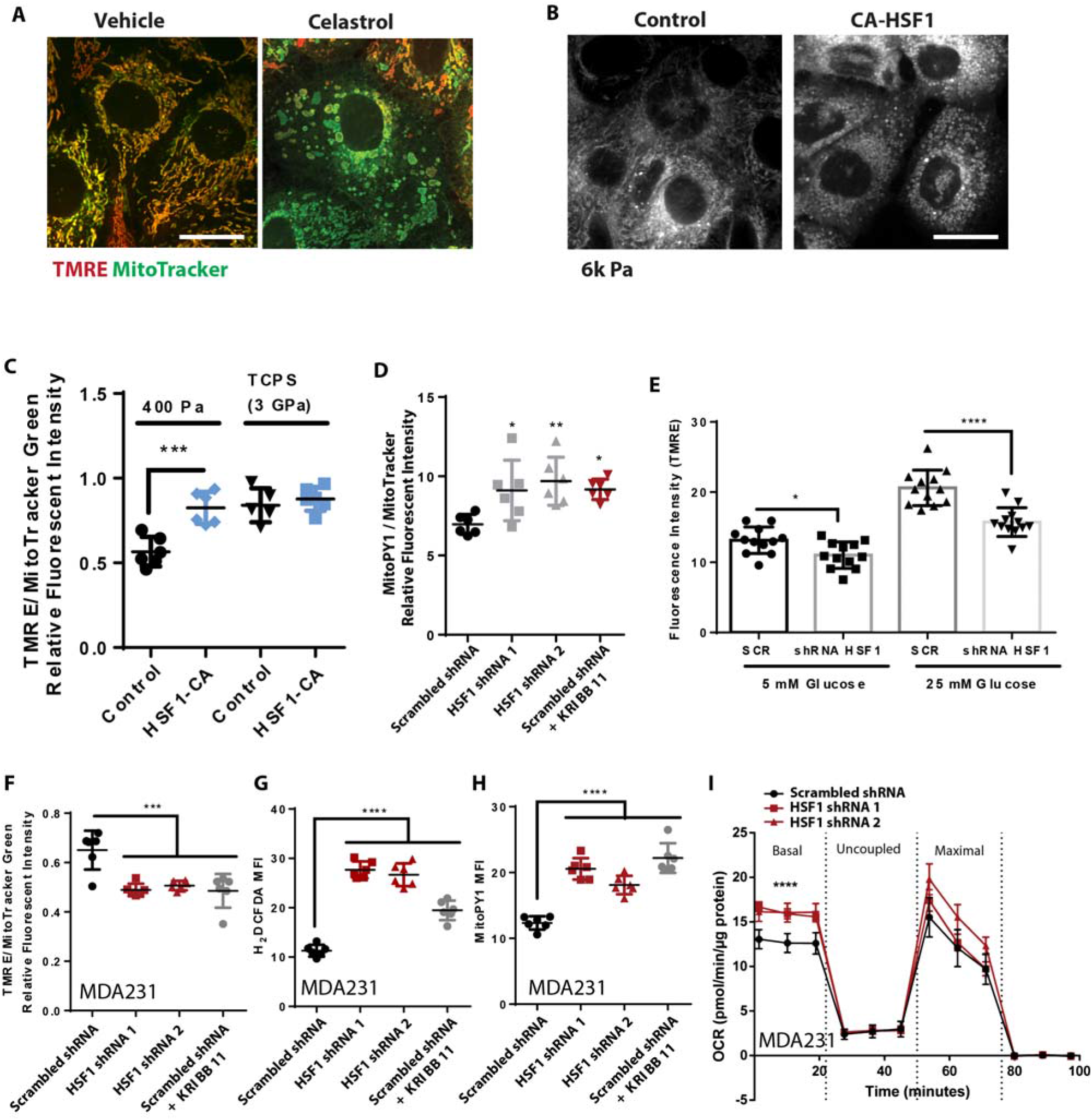
A. Representative microscopy depicting morphology and mitochondrial membrane potential staining of live cells via TMRE [10 nM] staining and mitotracker (green FM) [100 nM] +/- vehicle or Celastrol [2 μM] treatment for 40 minutes prior to imaging. (Scale Bar: 10 μm) B. Representative microscopy depicting mitochondrial network structure of cells cultured on 6k Pa ECM expressing constitutively active HSF1 or empty vector control, mitotracker (deep red FM) [100 nM]. (Scale Bar: 10 μm) C. Mitochondrial membrane potential MCF10A cells expressing CA-HSF1 on 400 Pa ECM or tissue culture polystyrene (TCPS) surfaces, measured with TMRE [10 nM] after 1 h staining (n=6) (n=6 wells, repeated 3 times). D. Mitochondrial hydrogen peroxide production of MCF10A cells expressing a scrambled shRNA +/- KRIBB11 [2 μM] or two different shRNAs targeting HSF1, measured with MitoPY1 [5 μM] normalized to Mitotracker green staining [100nM] (n=6 wells, repeated 3 times). E. Mitochondrial membrane potential of MDA-231 cells cultured on TCPS expressing a scrambled shRNA +/- shRNAs targeting HSF1 (HSF1 shRNA 1) +/- glucose [5 or 25 mM], measured with TMRE [10 nM] after 1 h staining (n=6 wells, repeated 3 times). F. Mitochondrial hydrogen peroxide production of MDA-231 cells cells expressing a scrambled shRNA +/- KRIBB11 [2 μM] or two different shRNAs targeting HSF1, measured with MitoPY1 [5 μM] normalized to Mitotracker green staining [100nM] (n=6 wells, repeated 3 times). G. Oxidative stress indicator intensity of of MDA-231 cells expressing a scrambled shRNA +/- KRIBB11 [2 μM] or two different shRNAs targeting HSF1, measured with H2DCFDA [2 μM] H. Mitochondrial membrane potential of MDA-231 cells expressing a scrambled shRNA +/- KRIBB11 [2 μM] or shRNAs targeting HSF1 +/- glucose [5 or 25 mM], measured with TMRE [10 nM] after 1 h staining (n=6 wells, repeated 3 times). I. OCR of MDA-231 cells expressing a scrambled shRNA or two different shRNAs targeting HSF1 (n=5 wells, 3 replicate measures, repeated 3 times) *Data shown represent ± SEM. *P < 0.05, **P < 0.01, ***P < 0.005, < 0.0001 via twotailed unpaired Student t test (E) or one-way ANOVA with Tukey test for multiple comparisons (B*).

**Figure S6.**
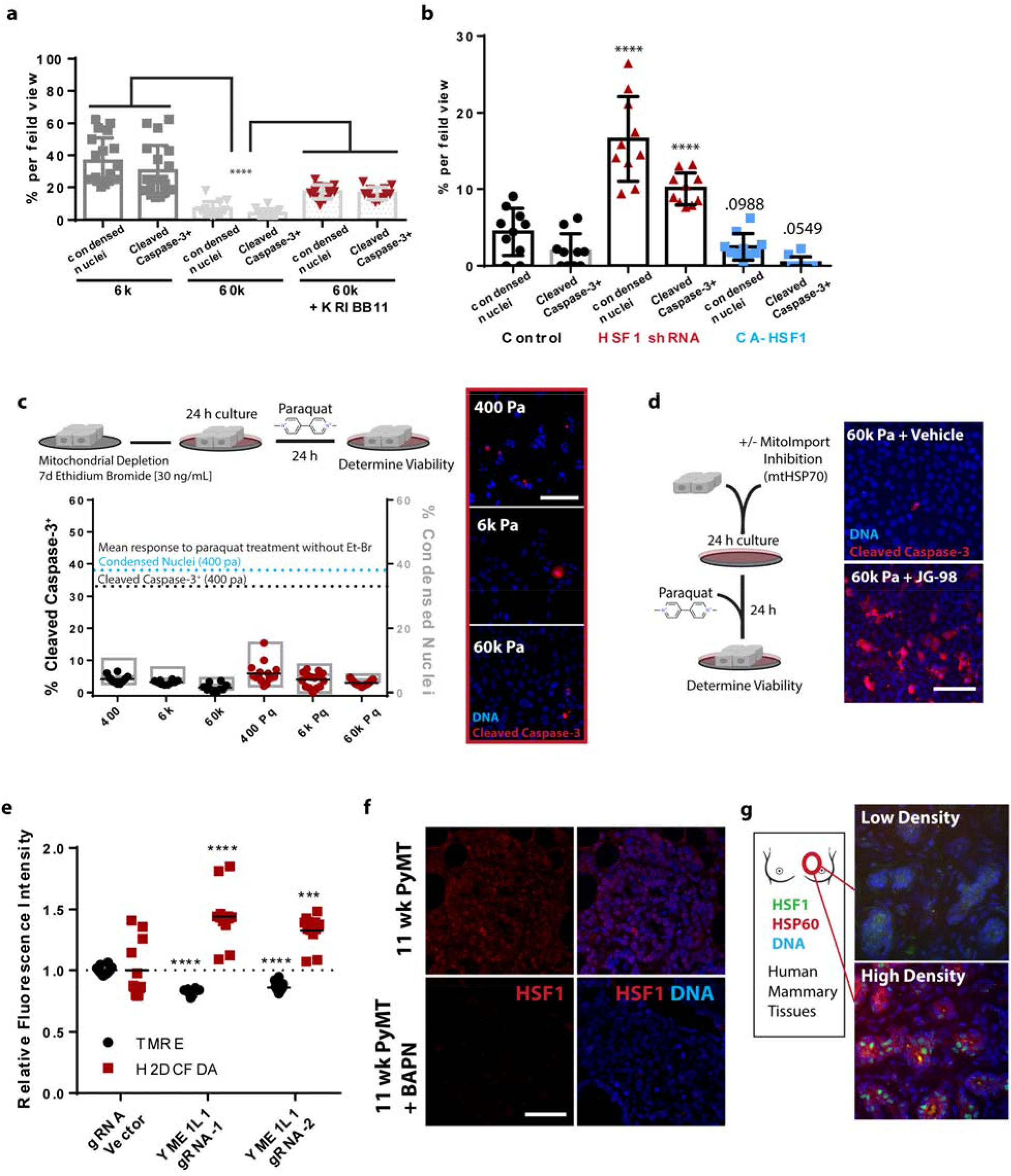
A. Cleaved caspase 3 staining (red) and nuclear condensation (dapi) of cells cultured on 6 or 60 Pa PA-gel surfaces for 24 h +/- vehicle or KRIBB11 [2 μM] followed by 24 h paraquat treatment [10 mM], (n=16 field views ~ 600 cells per condition, repeated 3 times) B. Quantitation of MCF10A cells expressing HSF1 shRNA 1 or CA-HSF1 from 10 field views for condensed nuclei and cleaved caspase-3 positive cells, cultured on fibronectin coated glass coverslips (~800 cells counted per condition, repeated 3 times). C. Representative microscopy and quantitation of indicators of apoptosis with cleaved caspase 3 staining (red) and nuclear condensation (dapi) of MCF10A cells previously cultured for 7 d with 30 ng/mL ethidium bromide in the culture media, then transferred and cultured on 400, 6k, 60k Pa PA-gel surfaces for 24 h with another 24 h +/- paraquat treatment [10 mM]. 100k cells/well of 24 well plate. EtBr maintained in media during paraquat challenge. (~600 cells counted per condition). Repeated 3 times (Scale Bar: 100 μm) D. Confocal microscopy of indicators of apoptosis with cleaved caspase 3 staining (red) and nuclear condensation (dapi) of cells cultured 60k Pa ECM surfaces for 24 h +/- JG98 [1 μM] followed by 24 h paraquat treatment [10 mM]. 100k cells/well of 24 well plate, repeated 2 times. E. Relative mitochondrial membrane potential and oxidative stress indicator intensity of MCF10A cells expressing YME1L1 knockdown via CRISPR-I compared to CRISPR-I and empty guide vector expressing cells on fibronectin coated TCPS, measured with TMRE [10 nM] and H_2_DCFDA [2 μM] for 1 h (n= 6 wells, repeated 2 times, both included) F. Representative immunofluorescence microscopy of murine mammary tumors derived from mice treated with BAPN, which inhibits lysyl oxidase mediated crosslinking of collagen and reduces ECM stiffness (*113*). Immunofluorescence microscopy for HSF1 (red) and DNA (blue). Supporting evidence showing that tumor HSF1 levels and nuclear localization may be sensitive to physical properties of the ECM. (Scale Bar: 100 μm). G. Representative immunofluorescence microscopy of human mammary glands derived from patient biopsies from low and high density breast tissues stained for HSF1 (green), HSP60 (red), and DNA (blue). In varied density human mammary tissues, HSF1 translocates to the nucleus and enhanced target gene expression in high density tissues supporting the in vitro evidence that HSF1 activity may be sensitive to ECM stiffness. (Scale Bar: 100 μm) *Data shown represent ± SEM. *P < 0.05, **P < 0.01, ***P < 0.005, < 0.0001 via twotailed unpaired Student t test (E) or one-way ANOVA with Tukey test for multiple comparisons (B)*.

**Supplemental Table 1.**
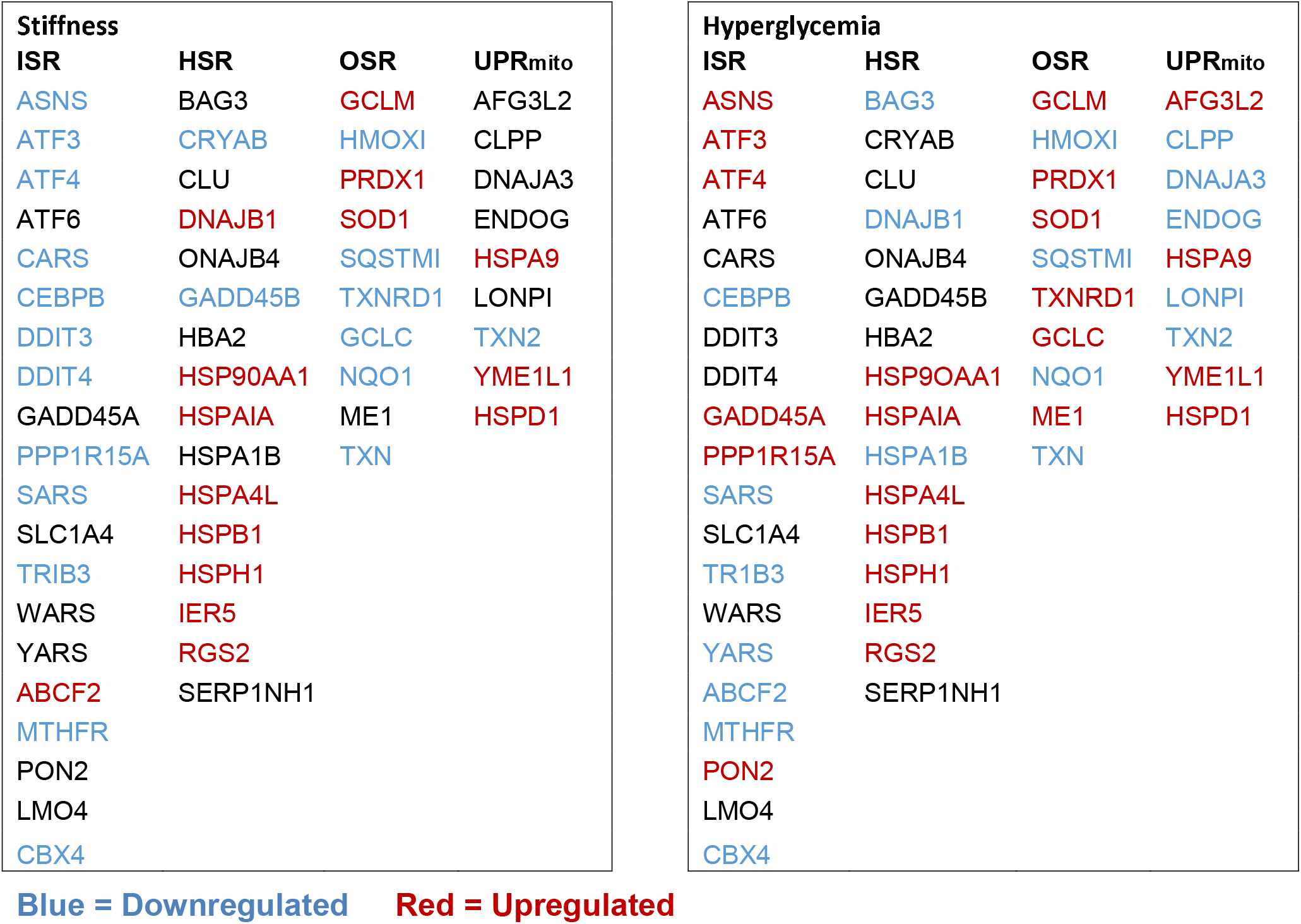

## References and Supplemental References

1. C. R. Reczek, N. S. Chandel, The Two Faces of Reactive Oxygen Species in Cancer. Annu. Rev. Cancer Biol. 1, 79–98 (2017).

2. D. C. Wallace, Mitochondria and cancer. Nat. Rev. Cancer. 12, 685–698 (2012).

3. F. Scialò, A. Sriram, D. Fernández-Ayala, N. Gubina, M. Lõhmus, G. Nelson, A. Logan, H. M. Cooper, P. Navas, J. A. Enríquez, M. P. Murphy, A. Sanz, Mitochondrial ROS Produced via Reverse Electron Transport Extend Animal Lifespan. Cell Metab. 23, 725–734 (2016).

4. R. S. Balaban, S. Nemoto, T. Finkel, Mitochondria, oxidants, and aging. Cell. 120, 483–495 (2005).

5. M. Fane, A. T. Weeraratna, How the ageing microenvironment influences tumour progression. Nat. Rev. Cancer. 20, 89–106 (2020).

6. S. Vyas, E. Zaganjor, M. C. Haigis, Mitochondria and Cancer. Cell. 166, 555–566 (2016).

7. W. Ladiges, J. Wanagat, B. Preston, L. Loeb, P. Rabinovitch, A mitochondrial view of aging, reactive oxygen species and metastatic cancer. Aging Cell. 9, 462–465 (2010).

8. N. Sun, R. J. Youle, T. Finkel, The Mitochondrial Basis of Aging. Mol. Cell. 61, 654–666 (2016).

9. Y. A. Miroshnikova, J. K. Mouw, J. M. Barnes, M. W. Pickup, J. N. Lakins, Y. Kim, K. Lobo, A. I. Persson, G. F. Reis, T. R. McKnight, E. C. Holland, J. J. Phillips, V. M. Weaver, Tissue mechanics promote IDH1-dependent HIF1α-tenascin C feedback to regulate glioblastoma aggression. Nat. Cell Biol. 18, 1336–1345 (2016).

10. M. J. Oudin, V. M. Weaver, Physical and Chemical Gradients in the Tumor Microenvironment Regulate Tumor Cell Invasion, Migration, and Metastasis. Cold Spring Harb. Symp. Quant. Biol. 81, 189–205 (2016).

11. J. M. Northcott, I. S. Dean, J. K. Mouw, V. M. Weaver, Feeling Stress: The Mechanics of Cancer Progression and Aggression. Front. cell Dev. Biol. 6, 17 (2018).

12. P. Schedin, P. J. Keely, Mammary gland ECM remodeling, stiffness, and mechanosignaling in normal development and tumor progression. Cold Spring Harb. Perspect. Biol. 3, a003228 (2011).

13. V. Anesti, L. Scorrano, The relationship between mitochondrial shape and function and the cytoskeleton. Biochim. Biophys. Acta. 1757, 692–699 (2006).

14. K. M. Tharp, M. S. Kang, G. A. Timblin, J. Dempersmier, G. E. Dempsey, P.-J. H. Zushin, J. Benavides, C. Choi, C. X. Li, A. K. Jha, S. Kajimura, K. E. Healy, H. S. Sul, K. Saijo, S. Kumar, A. Stahl, Actomyosin-Mediated Tension Orchestrates Uncoupled Respiration in Adipose Tissues. Cell Metab. 27, 602–615.e4 (2018).

15. M. J. Paszek, N. Zahir, K. R. Johnson, J. N. Lakins, G. I. Rozenberg, A. Gefen, C. A. Reinhart-King, S. S. Margulies, M. Dembo, D. Boettiger, D. A. Hammer, V. M. Weaver, Tensional homeostasis and the malignant phenotype. Cancer Cell. 8, 241–254 (2005).

16. R. Oria, T. Wiegand, J. Escribano, A. Elosegui-Artola, J. J. Uriarte, C. Moreno-Pulido, I. Platzman, P. Delcanale, L. Albertazzi, D. Navajas, X. Trepat, J. M. García-Aznar, E. A. Cavalcanti-Adam, P. Roca-Cusachs, Force loading explains spatial sensing of ligands by cells. Nature. 552, 219–224 (2017).

17. G. L. Lin, D. M. Cohen, R. A. Desai, M. T. Breckenridge, L. Gao, M. J. Humphries, C. S. Chen, Activation of beta 1 but not beta 3 integrin increases cell traction forces. FEBS Lett. 587, 763–769 (2013).

18. S. R. Caliari, J. A. Burdick, A practical guide to hydrogels for cell culture. Nat. Methods. 13, 405–414 (2016).

19. K. M. Tharp, V. M. Weaver, Modeling Tissue Polarity in Context. J. Mol. Biol. 430, 3613–3628 (2018).

20. D. T. Butcher, T. Alliston, V. M. Weaver, A tense situation: forcing tumour progression. Nat. Rev. Cancer. 9, 108–122 (2009).

21. D. R. Croft, M. F. Olson, Conditional regulation of a ROCK-estrogen receptor fusion protein. Methods Enzymol. 406, 541–553 (2006).

22. C. Yik-Sham Chung, G. A. Timblin, K. Saijo, C. J. Chang, Versatile Histochemical Approach to Detection of Hydrogen Peroxide in Cells and Tissues Based on Puromycin Staining. J. Am. Chem. Soc. 140, 6109–6121 (2018).

23. M. Sciacovelli, E. Gonçalves, T. I. Johnson, V. R. Zecchini, A. S. H. da Costa, E. Gaude, A. V. Drubbel, S. J. Theobald, S. R. Abbo, M. G. B. Tran, V. Rajeeve, S. Cardaci, S. Foster, H. Yun, P. Cutillas, A. Warren, V. Gnanapragasam, E. Gottlieb, K. Franze, B. Huntly, E. R. Maher, P. H. Maxwell, J. Saez-Rodriguez, C. Frezza, Fumarate is an epigenetic modifier that elicits epithelial-to-mesenchymal transition. Nature. 537, 544–547 (2016).

24. A. Barasa, G. Godina, P. Buffa, I. Pasquali-Ronchetti, Biochemical lesions of respiratory enzymes and configurational changes of mitochondria in vivo. I. The effect of fluoroacetate: a study by phase-contrast microscopy and time-lapse cinemicrography. Z. Zellforsch. Mikrosk. Anat. 138, 187–210 (1973).

25. D. Jimenez-Blasco, A. Busquets-Garcia, E. Hebert-Chatelain, R. Serrat, C. Vicente-Gutierrez, C. Ioannidou, P. Gómez-Sotres, I. Lopez-Fabuel, M. Resch-Beusher, E. Resel, D. Arnouil, D. Saraswat, M. Varilh, A. Cannich, F. Julio-Kalajzic, I. Bonilla-Del Río, A. Almeida, N. Puente, S. Achicallende, M.-L. Lopez-Rodriguez, C. Jollé, N. Déglon, L. Pellerin, C. Josephine, G. Bonvento, A. Panatier, B. Lutz, P.-V. Piazza, M. Guzmán, L. Bellocchio, A.-K. Bouzier-Sore, P. Grandes, J. P. Bolaños, G. Marsicano, Glucose metabolism links astroglial mitochondria to cannabinoid effects. Nature. 583, 603–608 (2020).

26. P. Lindström, J. Sehlin, Effect of glucose on the intracellular pH of pancreatic islet cells. Biochem. J. 218, 887–892 (1984).

27. Q. Wang, M. Zhang, G. Torres, S. Wu, C. Ouyang, Z. Xie, M.-H. Zou, Metformin Suppresses Diabetes-Accelerated Atherosclerosis via the Inhibition of Drp1-Mediated Mitochondrial Fission. Diabetes. 66, 193–205 (2017).

28. B.-C. Chen, W. R. Legant, K. Wang, L. Shao, D. E. Milkie, M. W. Davidson, C. Janetopoulos, X. S. Wu, J. A. Hammer 3rd, Z. Liu, B. P. English, Y. Mimori-Kiyosue, D. P. Romero, A. T. Ritter, J. Lippincott-Schwartz, L. Fritz-Laylin, R. D. Mullins, D. M. Mitchell, J. N. Bembenek, A.-C. Reymann, R. Böhme, S. W. Grill, J. T. Wang, G. Seydoux, U. S. Tulu, D. P. Kiehart, E. Betzig, Lattice light-sheet microscopy: imaging molecules to embryos at high spatiotemporal resolution. Science. 346, 1257998 (2014).

29. J. E. Aldridge, T. Horibe, N. J. Hoogenraad, Discovery of genes activated by the mitochondrial unfolded protein response (mtUPR) and cognate promoter elements. PLoS One. 2, e874–e874 (2007).

30. Y.-F. Lin, C. M. Haynes, Metabolism and the UPR(mt). Mol. Cell. 61, 677–682 (2016).

31. H.-G. Sprenger, T. Langer, The Good and the Bad of Mitochondrial Breakups. Trends Cell Biol. 29, 888–900 (2019).

32. R. J. Youle, A. M. van der Bliek, Mitochondrial fission, fusion, and stress. Science. 337, 1062–1065 (2012).

33. G. Twig, O. S. Shirihai, The interplay between mitochondrial dynamics and mitophagy. Antioxid. Redox Signal. 14, 1939–1951 (2011).

34. Y. Miyazono, S. Hirashima, N. Ishihara, J. Kusukawa, K.-I. Nakamura, K. Ohta, Uncoupled mitochondria quickly shorten along their long axis to form indented spheroids, instead of rings, in a fission-independent manner. Sci. Rep. 8, 350 (2018).

35. X. Liu, G. Hajnóczky, Altered fusion dynamics underlie unique morphological changes in mitochondria during hypoxia-reoxygenation stress. Cell Death Differ. 18, 1561–1572 (2011).

36. S. W. Perry, J. P. Norman, J. Barbieri, E. B. Brown, H. A. Gelbard, Mitochondrial membrane potential probes and the proton gradient: a practical usage guide. Biotechniques. 50, 98–115 (2011).

37. C.-H. Choi, B. A. Webb, M. S. Chimenti, M. P. Jacobson, D. L. Barber, pH sensing by FAK-His58 regulates focal adhesion remodeling. J. Cell Biol. 202, 849–859 (2013).

38. T. Tominaga, D. L. Barber, Na-H exchange acts downstream of RhoA to regulate integrin-induced cell adhesion and spreading. Mol. Biol. Cell. 9, 2287–2303 (1998).

39. T. Tominaga, T. Ishizaki, S. Narumiya, D. L. Barber, p160ROCK mediates RhoA activation of Na-H exchange. EMBO J. 17, 4712–4722 (1998).

40. K. Nehrke, J. E. Melvin, The NHX family of Na+-H+ exchangers in Caenorhabditis elegans. J. Biol. Chem. 277, 29036–29044 (2002).

41. M. Ristow, S. Schmeisser, Extending life span by increasing oxidative stress. Free Radic. Biol. Med. 51, 327–336 (2011).

42. P. S. Brookes, Y. Yoon, J. L. Robotham, M. W. Anders, S.-S. Sheu, Calcium, ATP, and ROS: a mitochondrial love-hate triangle. Am. J. Physiol. Cell Physiol. 287, C817–C833 (2004).

43. C. Giorgi, S. Marchi, P. Pinton, The machineries, regulation and cellular functions of mitochondrial calcium. Nat. Rev. Mol. Cell Biol. 19, 713–730 (2018).

44. P. Hernansanz-Agustín, C. Choya-Foces, S. Carregal-Romero, E. Ramos, T. Oliva, T. Villa-Piña, L. Moreno, A. Izquierdo-Álvarez, J. D. Cabrera-García, A. Cortés, A. V. Lechuga-Vieco, P. Jadiya, E. Navarro, E. Parada, A. Palomino-Antolín, D. Tello, R. Acín-Pérez, J. C. Rodríguez-Aguilera, P. Navas, Á. Cogolludo, I. López-Montero, Á. Martínez-Del-Pozo, J. Egea, M. G. López, J. W. Elrod, J. Ruíz-Cabello, A. Bogdanova, J. A. Enríquez, A. Martínez-Ruiz, Na(+) controls hypoxic signalling by the mitochondrial respiratory chain. Nature (2020), doi:10.1038/s41586-020-2551-y.

45. P. R. Castello, D. A. Drechsel, M. Patel, Mitochondria are a major source of paraquat-induced reactive oxygen species production in the brain. J. Biol. Chem. 282, 14186–14193 (2007).

46. J. Yun, T. Finkel, Mitohormesis. Cell Metab. 19, 757–766 (2014).

47. Q. Ma, Role of nrf2 in oxidative stress and toxicity. Annu. Rev. Pharmacol. Toxicol. 53, 401–426 (2013).

48. M. Ristow, Unraveling the truth about antioxidants: mitohormesis explains ROS-induced health benefits. Nat. Med. 20, 709–711 (2014).

49. M. P. Mattson, Hormesis defined. Ageing Res. Rev. 7, 1–7 (2008).

50. S. E. Calvo, K. R. Clauser, V. K. Mootha, MitoCarta2.0: an updated inventory of mammalian mitochondrial proteins. Nucleic Acids Res. 44, D1251–7 (2016).

51. J. Labbadia, R. M. Brielmann, M. F. Neto, Y.-F. Lin, C. M. Haynes, R. I. Morimoto, Mitochondrial Stress Restores the Heat Shock Response and Prevents Proteostasis Collapse during Aging. Cell Rep. 21, 1481–1494 (2017).

52. J. M. D. Grandjean, L. Plate, R. I. Morimoto, M. J. Bollong, E. T. Powers, R. L. Wiseman, Deconvoluting Stress-Responsive Proteostasis Signaling Pathways for Pharmacologic Activation Using Targeted RNA Sequencing. ACS Chem. Biol. 14, 784–795 (2019).

53. D. Reichmann, W. Voth, U. Jakob, Maintaining a Healthy Proteome during Oxidative Stress. Mol. Cell. 69, 203–213 (2018).

54. A. M. Nargund, M. W. Pellegrino, C. J. Fiorese, B. M. Baker, C. M. Haynes, Mitochondrial import efficiency of ATFS-1 regulates mitochondrial UPR activation. Science. 337, 587–590 (2012).

55. C. J. Fiorese, A. M. Schulz, Y.-F. Lin, N. Rosin, M. W. Pellegrino, C. M. Haynes, The Transcription Factor ATF5 Mediates a Mammalian Mitochondrial UPR. Curr. Biol. 26, 2037–2043 (2016).

56. A. Katiyar, M. Fujimoto, K. Tan, A. Kurashima, P. Srivastava, M. Okada, R. Takii, A. Nakai, HSF1 is required for induction of mitochondrial chaperones during the mitochondrial unfolded protein response. FEBS Open Bio (2020), doi:10.1002/2211-5463.12863.

57. F. Boos, L. Krämer, C. Groh, F. Jung, P. Haberkant, F. Stein, F. Wollweber, A. Gackstatter, E. Zöller, M. van der Laan, M. M. Savitski, V. Benes, J. M. Herrmann, Mitochondrial protein-induced stress triggers a global adaptive transcriptional programme. Nat. Cell Biol. 21, 442–451 (2019).

58. S.-G. Ahn, D. J. Thiele, Redox regulation of mammalian heat shock factor 1 is essential for Hsp gene activation and protection from stress. Genes Dev. 17, 516–528 (2003).

59. S. Paul, S. Ghosh, S. Mandal, S. Sau, M. Pal, NRF2 transcriptionally activates the heat shock factor 1 promoter under oxidative stress and affects survival and migration potential of MCF7 cells. J. Biol. Chem. 293, 19303–19316 (2018).

60. N. Wiedemann, N. Pfanner, Mitochondrial Machineries for Protein Import and Assembly. Annu. Rev. Biochem. 86, 685–714 (2017).

61. A. E. Charos, B. D. Reed, D. Raha, A. M. Szekely, S. M. Weissman, M. Snyder, A highly integrated and complex PPARGC1A transcription factor binding network in HepG2 cells. Genome Res. 22, 1668–1679 (2012).

62. M. L. Mendillo, S. Santagata, M. Koeva, G. W. Bell, R. Hu, R. M. Tamimi, E. Fraenkel, T. A. Ince, L. Whitesell, S. Lindquist, HSF1 drives a transcriptional program distinct from heat shock to support highly malignant human cancers. Cell. 150, 549–562 (2012).

63. S. E. LeBoeuf, W. L. Wu, T. R. Karakousi, B. Karadal, S. R. Jackson, S. M. Davidson, K.-K. Wong, S. B. Koralov, V. I. Sayin, T. Papagiannakopoulos, Activation of Oxidative Stress Response in Cancer Generates a Druggable Dependency on Exogenous Nonessential Amino Acids. Cell Metab. 31, 339–350.e4 (2020).

64. X. Ma, L. Xu, A. T. Alberobello, O. Gavrilova, A. Bagattin, M. Skarulis, J. Liu, T. Finkel, E. Mueller, Celastrol Protects against Obesity and Metabolic Dysfunction through Activation of a HSF1-PGC1α Transcriptional Axis. Cell Metab. 22, 695–708 (2015).

65. C. R. Reczek, K. Birsoy, H. Kong, I. Martínez-Reyes, T. Wang, P. Gao, D. M. Sabatini, N. S. Chandel, A CRISPR screen identifies a pathway required for paraquat-induced cell death. Nat. Chem. Biol. 13, 1274–1279 (2017).

66. O. Schmidt, N. Pfanner, C. Meisinger, Mitochondrial protein import: from proteomics to functional mechanisms. Nat. Rev. Mol. Cell Biol. 11, 655–667 (2010).

67. H. C. Schneider, J. Berthold, M. F. Bauer, K. Dietmeier, B. Guiard, M. Brunner, W. Neupert, Mitochondrial Hsp70/MIM44 complex facilitates protein import. Nature. 371, 768–774 (1994).

68. C. C. Deocaris, S. C. Kaul, R. Wadhwa, On the brotherhood of the mitochondrial chaperones mortalin and heat shock protein 60. Cell Stress Chaperones. 11, 116–128 (2006).

69. X. Li, S. R. Srinivasan, J. Connarn, A. Ahmad, Z. T. Young, A. M. Kabza, E. R. P. Zuiderweg, D. Sun, J. E. Gestwicki, Analogs of the Allosteric Heat Shock Protein 70 (Hsp70) Inhibitor, MKT-077, as Anti-Cancer Agents. ACS Med. Chem. Lett. 4, 1042–1047 (2013).

70. S. R. Srinivasan, L. C. Cesa, X. Li, O. Julien, M. Zhuang, H. Shao, J. Chung, I. Maillard, J. A. Wells, C. S. Duckett, J. E. Gestwicki, Heat Shock Protein 70 (Hsp70) Suppresses RIP1-Dependent Apoptotic and Necroptotic Cascades. Mol. Cancer Res. 16, 58–68 (2018).

71. T. MacVicar, T. Langer, OPA1 processing in cell death and disease - the long and short of it. J. Cell Sci. 129, 2297–2306 (2016).

72. T. MacVicar, Y. Ohba, H. Nolte, F. C. Mayer, T. Tatsuta, H.-G. Sprenger, B. Lindner, Y. Zhao, J. Li, C. Bruns, M. Krüger, M. Habich, J. Riemer, R. Schwarzer, M. Pasparakis, S. Henschke, J. C. Brüning, N. Zamboni, T. Langer, Lipid signalling drives proteolytic rewiring of mitochondria by YME1L. Nature. 575, 361–365 (2019).

73. J. R. Cantor, M. Abu-Remaileh, N. Kanarek, E. Freinkman, X. Gao, A. Louissaint Jr, C. A. Lewis, D. M. Sabatini, Physiologic Medium Rewires Cellular Metabolism and Reveals Uric Acid as an Endogenous Inhibitor of UMP Synthase. Cell. 169, 258–272.e17 (2017).

74. P. DelNero, B. D. Hopkins, L. C. Cantley, C. Fischbach, Cancer metabolism gets physical. Sci. Transl. Med. 10, eaaq1011 (2018).

75. Z. T. Schafer, A. R. Grassian, L. Song, Z. Jiang, Z. Gerhart-Hines, H. Y. Irie, S. Gao, P. Puigserver, J. S. Brugge, Antioxidant and oncogene rescue of metabolic defects caused by loss of matrix attachment. Nature. 461, 109–113 (2009).

76. E. Werner, Z. Werb, Integrins engage mitochondrial function for signal transduction by a mechanism dependent on Rho GTPases. J. Cell Biol. 158, 357–368 (2002).

77. D. C. Radisky, D. D. Levy, L. E. Littlepage, H. Liu, C. M. Nelson, J. E. Fata, D. Leake, E. L. Godden, D. G. Albertson, M. A. Nieto, Z. Werb, M. J. Bissell, Rac1b and reactive oxygen species mediate MMP-3-induced EMT and genomic instability. Nature. 436, 123–127 (2005).

78. K. A. Dill, Dominant forces in protein folding. Biochemistry. 29, 7133–7155 (1990).

79. M. Guo, A. F. Pegoraro, A. Mao, E. H. Zhou, P. R. Arany, Y. Han, D. T. Burnette, M. H. Jensen, K. E. Kasza, J. R. Moore, F. C. Mackintosh, J. J. Fredberg, D. J. Mooney, J. Lippincott-Schwartz, D. A. Weitz, Cell volume change through water efflux impacts cell stiffness and stem cell fate. Proc. Natl. Acad. Sci. U. S. A. 114, E8618–E8627 (2017).

80. K. A. Dill, J. L. MacCallum, The protein-folding problem, 50 years on. Science. 338, 1042–1046 (2012).

81. R. Higuchi-Sanabria, P. A. Frankino, J. W. Paul 3rd, S. U. Tronnes, A. Dillin, A Futile Battle? Protein Quality Control and the Stress of Aging. Dev. Cell. 44, 139–163 (2018).

82. S. C. J. Helle, Q. Feng, M. J. Aebersold, L. Hirt, R. R. Grüter, A. Vahid, A. Sirianni, S. Mostowy, J. G. Snedeker, A. Šarić, T. Idema, T. Zambelli, B. Kornmann, Mechanical force induces mitochondrial fission. Elife. 6, e30292 (2017).

83. B. A. Hassell, G. Goyal, E. Lee, A. Sontheimer-Phelps, O. Levy, C. S. Chen, D. E. Ingber, Human Organ Chip Models Recapitulate Orthotopic Lung Cancer Growth, Therapeutic Responses, and Tumor Dormancy In Vitro. Cell Rep. 21, 508–516 (2017).

84. J. Irianto, C. R. Pfeifer, R. R. Bennett, Y. Xia, I. L. Ivanovska, A. J. Liu, R. A. Greenberg, D. E. Discher, Nuclear constriction segregates mobile nuclear proteins away from chromatin. Mol. Biol. Cell. 27, 4011–4020 (2016).

85. P. Isermann, J. Lammerding, Consequences of a tight squeeze: Nuclear envelope rupture and repair. Nucleus. 8, 268–274 (2017).

86. M. S. Samuel, J. I. Lopez, E. J. McGhee, D. R. Croft, D. Strachan, P. Timpson, J. Munro, E. Schröder, J. Zhou, V. G. Brunton, N. Barker, H. Clevers, O. J. Sansom, K. I. Anderson, V. M. Weaver, M. F. Olson, Actomyosin-mediated cellular tension drives increased tissue stiffness and β-catenin activation to induce epidermal hyperplasia and tumor growth. Cancer Cell. 19, 776–791 (2011).

87. S. Santagata, R. Hu, N. U. Lin, M. L. Mendillo, L. C. Collins, S. E. Hankinson, S. J. Schnitt, L. Whitesell, R. M. Tamimi, S. Lindquist, T. A. Ince, High levels of nuclear heatshock factor 1 (HSF1) are associated with poor prognosis in breast cancer. Proc. Natl. Acad. Sci. U. S. A. 108, 18378–18383 (2011).

88. R. Scherz-Shouval, S. Santagata, M. L. Mendillo, L. M. Sholl, I. Ben-Aharon, A. H. Beck, D. Dias-Santagata, M. Koeva, S. M. Stemmer, L. Whitesell, S. Lindquist, The reprogramming of tumor stroma by HSF1 is a potent enabler of malignancy. Cell. 158, 564–578 (2014).

89. I. Acerbi, L. Cassereau, I. Dean, Q. Shi, A. Au, C. Park, Y. Y. Chen, J. Liphardt, E. S. Hwang, V. M. Weaver, Human breast cancer invasion and aggression correlates with ECM stiffening and immune cell infiltration. Integr. Biol. (Camb). 7, 1120–1134 (2015).

90. C. Dai, S. B. Sampson, HSF1: Guardian of Proteostasis in Cancer. Trends Cell Biol. 26, 17–28 (2016).

91. O. Y. Wouters, D. T. A. Ploeger, S. M. van Putten, R. A. Bank, 3,4-Dihydroxy-L-Phenylalanine as a Novel Covalent Linker of Extracellular Matrix Proteins to Polyacrylamide Hydrogels with a Tunable Stiffness. Tissue Eng. Part C. Methods. 22, 91–101 (2016).

92. S. Urlinger, U. Baron, M. Thellmann, M. T. Hasan, H. Bujard, W. Hillen, Exploring the sequence space for tetracycline-dependent transcriptional activators: novel mutations yield expanded range and sensitivity. Proc. Natl. Acad. Sci. U. S. A. 97, 7963–7968 (2000).

93. P. M. Quirós, M. A. Prado, N. Zamboni, D. D’Amico, R. W. Williams, D. Finley, S. P. Gygi, J. Auwerx, Multi-omics analysis identifies ATF4 as a key regulator of the mitochondrial stress response in mammals. J. Cell Biol. 216, 2027–2045 (2017).

94. A. Nakai, M. Suzuki, M. Tanabe, Arrest of spermatogenesis in mice expressing an active heat shock transcription factor 1. EMBO J. 19, 1545–1554 (2000).

95. J. Pouysségur, C. Sardet, A. Franchi, G. L’Allemain, S. Paris, A specific mutation abolishing Na+/H+ antiport activity in hamster fibroblasts precludes growth at neutral and acidic pH. Proc. Natl. Acad. Sci. U. S. A. 81, 4833–4837 (1984).

96. N. A. Baird, P. M. Douglas, M. S. Simic, A. R. Grant, J. J. Moresco, S. C. Wolff, J. R. Yates 3rd, G. Manning, A. Dillin, HSF-1-mediated cytoskeletal integrity determines thermotolerance and life span. Science. 346, 360–363 (2014).

97. T. Yoneda, C. Benedetti, F. Urano, S. G. Clark, H. P. Harding, D. Ron, Compartment-specific perturbation of protein handling activates genes encoding mitochondrial chaperones. J. Cell Sci. 117, 4055–4066 (2004).

98. J. R. Daniele, D. J. Esping, G. Garcia, L. S. Parsons, E. A. Arriaga, A. Dillin, “High-Throughput Characterization of Region-Specific Mitochondrial Function and Morphology.” Sci. Rep. 7, 6749 (2017).

99. S. Liu, M. J. Mlodzianoski, Z. Hu, Y. Ren, K. McElmurry, D. M. Suter, F. Huang, sCMOS noise-correction algorithm for microscopy images. Nat. Methods. 14, 760–761 (2017).

100. T. (2019). Lambert, tlambert03/LLSpy: Lattice light-sheet post-processing utility. Zenodo (2019), doi:doi:10.5281/zenodo.1059099.

101. L. A. Royer, M. Weigert, U. Günther, N. Maghelli, F. Jug, I. F. Sbalzarini, E. W. Myers, ClearVolume: open-source live 3D visualization for light-sheet microscopy. Nat. Methods. 12, 480–481 (2015).

102. A. Dobin, C. A. Davis, F. Schlesinger, J. Drenkow, C. Zaleski, S. Jha, P. Batut, M. Chaisson, T. R. Gingeras, STAR: ultrafast universal RNA-seq aligner. Bioinformatics. 29, 15–21 (2013).

103. Y. C. Lin, S. Jhunjhunwala, C. Benner, S. Heinz, E. Welinder, R. Mansson, M. Sigvardsson, J. Hagman, C. A. Espinoza, J. Dutkowski, T. Ideker, C. K. Glass, C. Murre, A global network of transcription factors, involving E2A, EBF1 and Foxo1, that orchestrates B cell fate. Nat. Immunol. 11, 635–643 (2010).

104. M. B. Eisen, P. T. Spellman, P. O. Brown, D. Botstein, Cluster analysis and display of genome-wide expression patterns. Proc. Natl. Acad. Sci. U. S. A. 95, 14863–14868 (1998).

105. Y. Zhou, B. Zhou, L. Pache, M. Chang, A. H. Khodabakhshi, O. Tanaseichuk, C. Benner, S. K. Chanda, Metascape provides a biologist-oriented resource for the analysis of systems-level datasets. Nat. Commun. 10, 1523 (2019).

106. F. Meier, A.-D. Brunner, S. Koch, H. Koch, M. Lubeck, M. Krause, N. Goedecke, J. Decker, T. Kosinski, M. A. Park, N. Bache, O. Hoerning, J. Cox, O. Räther, M. Mann, Online Parallel Accumulation-Serial Fragmentation (PASEF) with a Novel Trapped Ion Mobility Mass Spectrometer. Mol. Cell. Proteomics. 17, 2534–2545 (2018).

107. J. Cox, M. Mann, MaxQuant enables high peptide identification rates, individualized p.p.b.-range mass accuracies and proteome-wide protein quantification. Nat. Biotechnol. 26, 1367–1372 (2008).

108. M. Choi, C.-Y. Chang, T. Clough, D. Broudy, T. Killeen, B. MacLean, O. Vitek, MSstats: an R package for statistical analysis of quantitative mass spectrometry-based proteomic experiments. Bioinformatics. 30, 2524–2526 (2014).

109. W. K. Russell, Z. Y. Park, D. H. Russell, Proteolysis in mixed organic-aqueous solvent systems: applications for peptide mass mapping using mass spectrometry. Anal. Chem. 73, 2682–2685 (2001).

110. D. Giustarini, I. Dalle-Donne, A. Milzani, P. Fanti, R. Rossi, Analysis of GSH and GSSG after derivatization with N-ethylmaleimide. Nat. Protoc. 8, 1660–1669 (2013).

111. D. I. Benjamin, A. Cozzo, X. Ji, L. S. Roberts, S. M. Louie, M. M. Mulvihill, K. Luo, D. K. Nomura, Ether lipid generating enzyme AGPS alters the balance of structural and signaling lipids to fuel cancer pathogenicity. Proc. Natl. Acad. Sci. U. S. A. 110, 14912–14917 (2013).

112. S. M. Louie, E. A. Grossman, L. A. Crawford, L. Ding, R. Camarda, T. R. Huffman, D. K. Miyamoto, A. Goga, E. Weerapana, D. K. Nomura, GSTP1 Is a Driver of Triple-Negative Breast Cancer Cell Metabolism and Pathogenicity. Cell Chem. Biol. 23, 567–578 (2016).

113. K. R. Levental, H. Yu, L. Kass, J. N. Lakins, M. Egeblad, J. T. Erler, S. F. T. Fong, K. Csiszar, A. Giaccia, W. Weninger, M. Yamauchi, D. L. Gasser, V. M. Weaver, Matrix crosslinking forces tumor progression by enhancing integrin signaling. Cell. 139, 891–906 (2009).

